# Annelid methylomes reveal ancestral developmental and ageing-associated epigenetic erosion across Bilateria

**DOI:** 10.1101/2023.12.21.572802

**Authors:** Kero Guynes, Luke A. Sarre, Allan M. Carrillo-Baltodano, Billie E. Davies, Lan Xu, Yan Liang, Francisco M. Martín-Zamora, Paul J. Hurd, Alex de Mendoza, José M. Martín-Durán

## Abstract

**Background:** DNA methylation in the form of 5-methylcytosine (5mC) is the most abundant base modification in animals. However, 5mC levels vary widely across taxa. While vertebrate genomes are hypermethylated, in most invertebrates, 5mC concentrates on constantly and highly transcribed genes (gene body methylation; GbM) and, in some species, on transposable elements (TEs), a pattern known as ‘mosaic’. Yet, the role and developmental dynamics of 5mC and how these explain interspecific differences in DNA methylation patterns remain poorly understood, especially in Spiralia, a large clade of invertebrates comprising nearly half of the animal phyla.

**Results:** Here, we generate base-resolution methylomes for three species with distinct genomic features and phylogenetic positions in Annelida, a major spiralian phylum. All possible 5mC patterns occur in annelids, from typical invertebrate intermediate levels in a mosaic distribution to hypermethylation and methylation loss. GbM is common to annelids with 5mC, and methylation differences across species are explained by taxon-specific transcriptional dynamics or the presence of intronic TEs. Notably, the link between GbM and transcription decays during development, and there is a gradual and global, age-dependent demethylation in adult stages. Moreover, reducing 5mC levels with cytidine analogues during early development impairs normal embryogenesis and reactivates TEs in the annelid *Owenia fusiformis*.

**Conclusions:** Our study indicates that global epigenetic erosion during development and ageing is an ancestral feature of bilateral animals. However, the tight link between transcription and gene body methylation is likely important in early embryonic stages, and 5mC-mediated TE silencing probably emerged convergently across animal lineages.

## Background

The reversible and heritable methylation of DNA, predominantly in the fifth position of the aromatic ring of a cytosine (i.e., 5-methylcytosine, or 5mC) of CpG dinucleotides, is a base modification that modulates diverse biological processes in animals and other eukaryotic lineages [1–3]. DNA methyltransferases (DNMTs) deposit this DNA modification, which is erased by ten-eleven translocation (TET) dioxygenases and bound by proteins such as methyl-CpG-binding domain (MBD) proteins [4–6]. The main DNMT families in animals include the maintenance type DNMT1 and the *de novo* DNMT3, whereas DNMT2 is a conserved tRNA methyltransferase [5]. Although this genetic toolkit to deposit, read, and erase 5mC is generally conserved in animals [7, 8], the genomic distribution of this base modification varies dramatically across the phyla in which it has been investigated [1, 2, 6]. Vertebrate genomes are hypermethylated as 5mC is widespread and occurs at high levels except at promoters and active distal regulatory elements [9]. In contrast, 5mC is sparsely distributed in most invertebrate genomes, exhibiting a mosaic pattern primarily concentrated in active gene bodies and sometimes in transposable elements (TEs) [10–12]. However, some invertebrates have secondarily diverged from this condition and display vertebrate-like hypermethylated [7] or, more frequently, unmethylated genomes [13, 14].

These contrasting genomic landscapes correlate with potentially distinct roles of 5mC in animal lineages. While DNA methylation is essential for normal embryogenesis in the hypermethylated vertebrate genomes [15–17], the function of this methyl mark is less understood in invertebrates because most of the work has focussed on lineages with generally low methylation levels, such as insects [18, 19], and the two best-established invertebrate systems, the fly *Drosophila melanogaster* and the nematode worm *Caenorhabditis elegans*, lack this epigenetic mark [13, 14]. Nonetheless, 5mC has been proposed to contribute to gene regulation in invertebrates by repressing intragenic spurious transcription start sites [20], modulating gene transcription levels in response to environmental cues [21, 22], and controlling the activity of *cis*-regulatory elements [23]. However, the “epigenetic reprogramming” occurring in early embryogenesis in vertebrates has not been observed in invertebrates [24], where developmental 5mC levels are generally constant, with recent reports of late 5mC developmental dynamics in invertebrate deuterostomes [23, 25]. The evolution and ancestral role of DNA methylation in animal genomes thus remains poorly understood, mainly because we lack comprehensive functional studies that profile 5mC at base resolution during development and across phylogeny for most invertebrate lineages.

Spiralia (also known as Lophotrochozoa) is one of the three major clades of bilaterally symmetrical animals, including species with economic, ecological, and societal importance [26]. Despite comprising nearly half of all animal phyla [27], our knowledge of genome regulation in this animal clade is limited [28, 29], which, together with the more divergent genomic features of traditional invertebrate models (e.g., insects and nematode worms), ultimately impacts our capacity to reconstruct ancestral characters for crucial nodes of the animal tree of life, most notably the last common bilaterian ancestor. The study of DNA methylation has mainly focused on only four of the 15 spiralian groups, namely rotifers, platyhelminthes, molluscs, and annelids, most often using *in-silico* predictions and low-resolution profiling techniques. Rotifers have a divergent DNA methylation toolkit, lacking DNMTs and TET genes [30, 31]. Instead of 5mC, they rely on a horizontally acquired bacterial methyltransferase that modifies their cytosines in the fourth position of the pyrimidine ring (4mC) to regulate TE activity [32]. 5mC DNA methylation is also absent in some platyhelminthes, such as the planarian *Schmidtea mediterranea* [33] and probably the parasite *Schistosoma mansoni* [34]. It is, however, present at low levels in *Macrostomum lignano* [35], where both DNMT1 and DNMT3 are conserved. Rotifers and platyhelminthes have fast molecular and genomic evolution rates, and thus, their divergent DNA methylation patterns are unlikely to represent the ancestral spiralian condition.

Molluscs and annelids are species-rich spiralian clades with more conservatively evolving genomes [28]. In molluscs, studies of DNA methylation have primarily focused on bivalves (e.g., the oyster *Crassostrea gigas*) and cephalopods [36–41], revealing generally low-to-moderate methylation levels (∼10% mCG). Base-resolution profiling in adult tissues with whole-genome bisulphite sequencing confirmed a mosaic 5mC landscape that concentrates in gene bodies of highly expressed genes and in young genic TEs of bivalves but not cephalopods [39, 41]. In contrast, some annelids display moderate to high (∼40–80%) methylation levels, such as nereidids (e.g., *Platynereis dumerilii* and *Alitta succinea*) and perhaps the deep-sea siboglinid *Riftia pachyptila* [8, 36], although these methylation levels are not substantiated with a reference genome and unbiased genome-wide data. Yet, other deep-sea annelids have lower global 5mC levels (∼20%) [42]. Genome-wide 5mC landscapes available for adult tissue of the deep-sea annelids support the enrichment of this epigenetic mark in actively transcribed gene bodies, but not TEs [42]. Notably, treatments with cytidine analogues aimed at inhibiting DNMTs impair molluscan development and annelid posterior regeneration [8, 40]. Therefore, despite their different methylation levels, molluscs and annelids display canonical mosaic methylation, unlike rotifers and most platyhelminthes. However, the limited taxonomic sampling, especially at critical nodes of the molluscan and annelid phylogeny, and the lack of temporal resolution for genome-wide methylomes during the life cycles of these animals prevent identifying how dynamic this epigenetic mark is, thereby hampering the reconstruction of DNA methylation evolution in Spiralia, and indeed Bilateria generally.

To address this gap, we comprehensively characterised the DNA methylation landscapes of three annelid species with distinct genomic features and phylogenetic positions within Annelida currently used as developmental model systems (Fig. 1a, b). *Owenia fusiformis* belongs to the sister clade to all remaining annelids [43], and its slow-evolving genome with an ancestral indirect larval development has helped to infer ancestral characters to Annelida [29, 44]. *Dimorphilus gyrociliatus* underwent morphological miniaturisation, has secondarily evolved direct development, and has one of the smallest known genomes for a free-living animal, almost devoid of TEs [45]. Finally, *Capitella teleta* has an indirect life cycle that can be closed in the lab and exhibits a slow-evolving genome [46, 47]. Using base pair resolution genome-wide profiling approaches, our findings demonstrate that 5mC levels vary during annelid development and across the annelid phylogeny, with *O. fusiformis* and *C. teleta* displaying moderate levels and a typical mosaic pattern but *D. gyrociliatus* showing negligible methylation as adults. DNA methylation positively correlates with transcriptional levels and stability, and normal methylation levels are essential for successful embryogenesis in *O. fusiformis*. However, the global DNA methylomes erode along development and aging in these annelids. Altogether, our data reveal a dynamic DNA methylation landscape in Annelida, indicating that age-dependent methylation around active gene bodies is an ancestral genome regulatory state for bilaterians, with recurrent transitions to hyper- and unmethylated states being more frequent than previously estimated.

**Figure 1.**
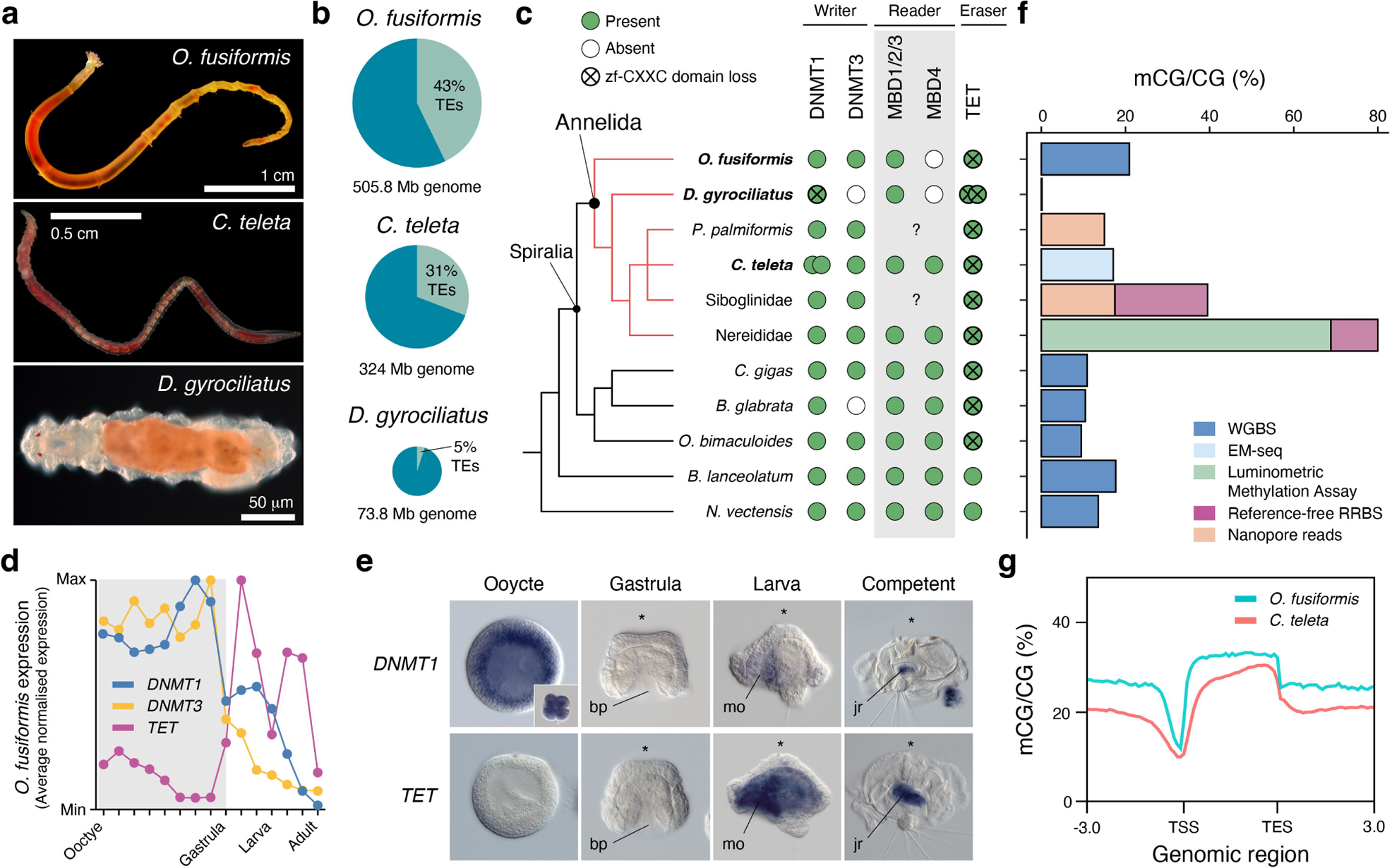
Annelids have different global DNA methylation levels. (**a**) Photographs of the three studied annelids. (**b**) Pie charts in scale representing genome size and percentage of repetitive elements in the genomes of *O. fusiformis*, *C. teleta* and *D. gyrociliatus*. (**c**) On the left, cladogram with the annelid lineages with existing global genome-wide 5mC and other representative metazoans. On the right, composition of the DNA methylation toolkit for the lineages on the left. (**f**) Bar plot representing the global methylation levels for representative annelid and animal lineages, colour-coded based on the methodology used. (**d**) Relative dynamics of writers (DNMT1 and DNMT3) and erasers (TET) of 5mC in the annelid *O. fusiformis*. Gastrulation is at the transition between an early phase dominated by DNA methylation writers and a later one when the expression of the eraser is predominant. (**e**) Whole-mount in situ hybridisation of the writer *DNMT1* and the eraser *TET* in the oocyte (a cleavage stage in the inset), and gastrula, larva and competent stages in *O. fusiformis*. *DNMT1* is expressed strongly in the oocyte and cleavage stages and more weakly in the ventral side of the larva and juvenile rudiment of the competent. Conversely, *TET* is not detected during early embryogenesis but more broadly in the larva and juvenile rudiment. (**g**) 5mC metagene profiles in *O. fusiformis* and *C. teleta*. These two annelids show canonical gene body methylation and 5mC depletion in promoters.

## Results

### The developmental dynamics of the DNA methylation toolkit in Annelida

To investigate the evolution of DNA methylation in Annelida, we first characterised the repertoire of genes involved in transferring methyl groups to cytosines in the DNA (DNMT1 and DNMT3 genes) and reading (MBD genes) and erasing (TET genes) this base modification in the genomes of *O. fusiformis*, *D. gyrociliatus*, and *C. teleta* (Fig. 1c; Fig. S1– 3). Consistent with previous results in the polychaete *P. dumerilii* and the leech *Helobdella robusta* [8], these three annelids have a DNMT1 gene, which in *C. teleta* is duplicated (Fig. S1). Notably, only the DNMT1 of *O. fusiformis* has a protein domain composition identical to the DNMT1 of humans (Fig. S4a). In *C. teleta* and *D. gyrociliatus*, the DNMT1 genes lack the most N-terminal DMAP domain (Fig. S4a). In addition, the DNMT1 of *D. gyrociliatus* and one of the paralogs of *C. teleta* lack the zinc finger CXXC domain, while the other paralog of *C. teleta* has this domain but not the DNMT1-RFD (Fig. S4a). However, despite these divergences in domain composition, the DNMT1 genes of these three annelids have conserved residues in the functionally active motifs of the methylase domain and could thus potentially methylate DNA (Fig. S4b). DNMT3 shows a conserved protein domain architecture in *O. fusiformis* and *C. teleta,* while it is absent in *D. gyrociliatus* (Fig. S1; Fig. S4c). Regarding the readers, the three species have an MBD1/2/3 gene (Fig. S2). Like *P. dumerilii* and *H. robusta* [8], *C. teleta* also has an MBD4 ortholog, absent in *O. fusiformis* and *D. gyrociliatus* (Fig. S2). In addition, all three annelids have a TET gene, duplicated in *D. gyrociliatus* (Fig. S3). Like many other invertebrates and the TET2 paralog of humans, the TET genes lack a zinc finger CXXC domain in annelids (Fig. 1c; Fig. S4e), suggesting ancestral or recurrent losses of this domain in Spiralia. Therefore, our findings support that a conserved DNA methylation machinery is an ancestral character of Annelida [8, 42]. Yet, this complement has diverged in specific annelid lineages, particularly regarding the structural domain composition of DNMTs.

The DNA methylation toolkit is dynamically expressed during the life cycles of *O. fusiformis*, *C. teleta*, and *D. gyrociliatus*. In these three species, DNMT1 transcripts (*DNMT1b* in *C. teleta*) are more abundant in early than late embryogenesis (Fig. S5a, c, d) and broadly mirror the transcriptional dynamics of *UHRF1*, the E3 ubiquitin ligase that recruits DNMT1 to hemimethylated DNA during S-phase [48, 49]. *DNMT3* genes are expressed at low levels (Fig. S5a, c), and the *MBD1/2/3* readers show a peak of expression during gastrulation and early embryogenesis (Fig. S5a, c, d). However, *TET* genes increase their expression right before gastrulation and peak at larval stages in *O. fusiformis* and *C. teleta* and the adults of *D. gyrociliatus* (for *TETa*) (Fig. S5a, c, d). Therefore, DNMT and TET genes exhibit inverse expression dynamics during annelid embryogenesis (Fig. 1d), as observed in some invertebrate deuterostomes [23].

To validate these temporal transcriptional dynamics, we characterise the spatial expression patterns of the DNA methylation toolkit during *O. fusiformis* embryogenesis. *DNMT1* is detected in the oocyte and fast-cycling blastomeres during early cleavage in this annelid (Fig. 1e; Fig. S5b). While no expression pattern was evident during gastrulation and late embryogenesis, *DNMT1* localised to the early larva’s ventral and foregut regions and the competent larva’s juvenile rudiment (Fig. 1e; Fig. S5b). No expression pattern was detected for the low-expressed *DNMT3* in *O. fusiformis* (Fig. S5b), whereas *MBD1/2/3* is ubiquitously detected during embryogenesis and later localises around the foregut, ventral side, and juvenile rudiment during early and late larval stages (Fig. S5b). Finally, *TET* first becomes diffusely expressed in the gastrula of *O. fusiformis* and is strongly detected in the gut and ventral region of the early larva and juvenile rudiment of the competent larva (Fig. 1e; Fig. S5b). DNA methylation writers (DNMT1 and DNMT3) and erasers (TET) thus localise to proliferative and differentiating tissues, suggesting a potential role of DNA methylation in regulating transcriptional programmes in this animal group.

### Annelida present the whole diversity of animal methylation patterns

To confirm the presence of DNA methylation in *O. fusiformis*, *D. gyrociliatus*, and *C. teleta*, we generated base pair resolution, genome-wide methylomes for the adults of these three species (Table S1). In their adult forms, *O. fusiformis* and *C. teleta* show 5mC in 20.92% and 17.1% of their CGs, respectively (Fig. 1f). However, *D. gyrociliatus* has negligible 5mC levels (0.16%), below the bisulfite non-conversion rate (Fig. 1f). Although this does not exclude the potential occurrence of DNA methylation at earlier stages of the life cycle in this miniature annelid, the ratio of observed vs expected genomic CpGs in *D. gyrociliatus* is close to equilibrium (0.9, Table S2) typical of unmethylated/lowly methylated species, suggesting lack of CG methylation despite encoding DNMT1 (Fig. S1; Fig. S4a). The DNA methylation levels observed in *O. fusiformis* and *C. teleta* coincide with those reported in other invertebrate lineages (10–20% CpG methylation levels), within (e.g., the molluscs *C. gigas*, *B. glabrata* and *O. bimaculoides*) and outside Spiralia (e.g., the lancelet *B. lanceolatum* or the sea anemone *N. vectensis*), but starkly contrast with those of some annelid lineages belonging to Nereididae, such as *P. dumerilii* and *Alitta succinea*, which have well over 60% of their CGs methylated (Fig. 1f) [8, 36], albeit the available data for these species is not genome-wide. Therefore, Annelida shows diverse levels of DNA methylation, from the possibly ancestral condition of low-to-moderate levels observed in *O. fusiformis* and *C. teleta* to the loss of this base modification in *D. gyrociliatus* and the hypermethylated state in some lineages.

### Global methylation levels decrease and decouple from transcription along development

Methylated CpGs show a mosaic distribution in *O. fusiformis* and *C. teleta*, concentrated within gene bodies (Fig. 1g; Fig. S6a, b). In both species, the promoter region upstream to the transcription start site (TSS) is hypomethylated, and there is between 1.1–1.2-fold increase in methylation levels in gene bodies compared with the background intergenic regions (Fig. 1g). To investigate whether the DNA methylation profiles observed in the adults of *O. fusiformis* and *C. teleta* were representative of their entire life cycles, we additionally generated whole-genome methylomes at one embryonic (gastrula, as the time point when DNMTs and TET change transcriptional dynamics; Fig. 1d) and one larval stage. In both annelids, there are higher 5mC levels in gastrulae (36.1% in *O. fusiformis* and 20.99% in *C. teleta*) and larvae (35.42% in *O.fusiformis* and 18.93% in *C. teleta*) than in adults (Fig. 2a, b). Despite these differences, the global pattern of CG methylation remains unchanged during the life cycle of these annelids, with a mosaic distribution concentrated in gene bodies and hypomethylated upstream promoters throughout (Fig. S6a, b, d, e). Therefore, the annelids *O. fusiformis* and *C. teleta* have typical invertebrate mosaic DNA methylation landscapes [1]. However, these erode as the life cycle progresses, as observed in two other phylogenetically distant invertebrates [23].

**Figure 2.**
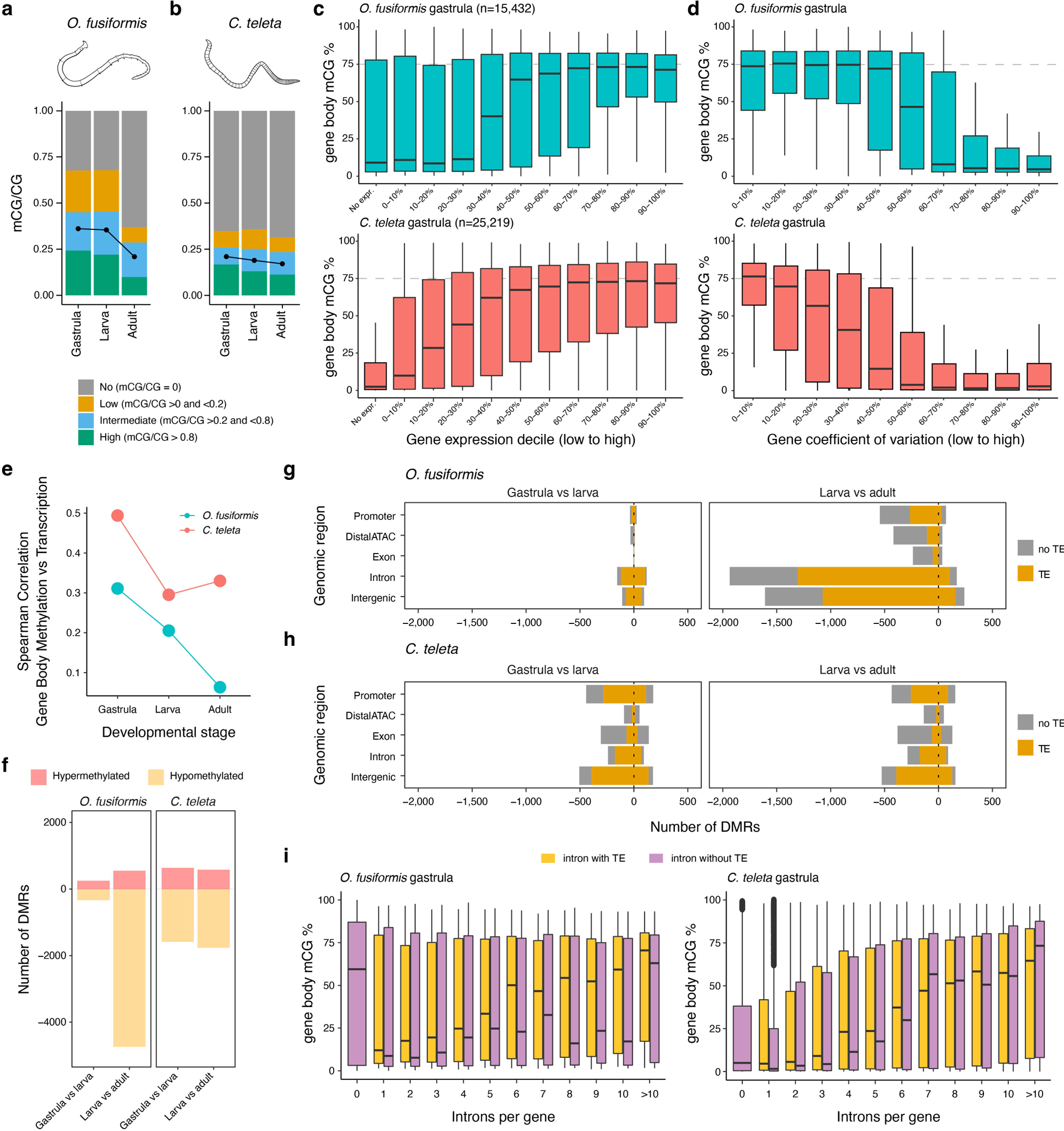
DNA methylation correlates with high and stable transcription. (**a**, **b**) Global CG methylation levels decrease during *O. fusiformis* (WGBS, **a**) and *C. teleta* (EM-seq, **b**) life cycles. Only CpGs with >10x coverage are depicted. (**c**) Box plots correlating gene body methylation and expression levels in *O. fusiformis* (top) and *C. teleta* (bottom) at the gastrula stage. Genes were divided into 10 deciles and non-expressed (TPM <1). (**d**) Box plots correlating gene body methylation and transcription stability in *O. fusiformis* (top) and *C. teleta* (bottom) at the gastrula stage. The coefficient of variation was calculated for all genes to rank them in 10 deciles from lower to higher values. Genes with a lower coefficient of variation (i.e., more transcriptionally stable) have higher gene body methylation levels in both species. (**e**) Correlation between gene body methylation and transcription in *O. fusiformis* (green line) and *C. teleta* (red line). As the life cycle progresses, gene body methylation and transcription are less strongly correlated. (**f**) Bar plots indicating the number of differentially methylated regions (DMRs) between pairwise comparisons during *O. fusiformis* (left) and *C. teleta* (right) life cycles. As expected by the gradual de-methylation, most DMRs represent a transition to a hypomethylated state. (**g**, **h**) Bar plots indicating the genomic annotation of the DMRs in *O. fusiformis* (**g**, top) and *C. teleta* (**h**, bottom). Most hypomethylated DMRs have a transposable element (TE) except in exonic and distal regulatory regions. (**i**) Box plots showing the effect of intronic transposable elements (TEs) in gene body methylation levels in *O. fusiformis* and *C. teleta* at the gastrula stage. Transcriptional units with an intronic TE show higher gene body methylation levels than equivalent genes without a TE in *O. fusiformis*, but not *C. teleta*.

To elucidate the interplay between gene body methylation (GbM) and gene expression, we first compared the GbM levels and transcriptional dynamics (transcriptional levels and stability) in the gastrula, larva and adult stages in *O. fusiformis* and *C. teleta*. Higher GbM correlates with higher gene expression values (Fig. 2c; Fig. S7a, c) and transcriptional stability (Fig. 2d; Fig. S7b, d) in both annelids, especially at the gastrula stage. Notably, genes in the highest expressed deciles tend to decrease their GbM levels more acutely than the rest as global methylation lowers during development (Fig. S7a, c) and, thereby, there is an overall decrease in the correlation between GbM and transcriptional levels as the life cycle progresses in both annelids, but especially in *O. fusiformis* (Fig. 2e). Therefore, DNA methylation and gene expression are positively correlated in annelids, although this association weakens as the life cycle progresses.

The drastic change in global methylation levels from larval to adult stages could be involved in regulating specific developmental processes. To examine this hypothesis, we first identified the genes whose GbM levels varied the most between each time point of development in *O. fusiformis* and *C. teleta* (Fig. S8a, c). 1,860 genes in *O. fusiformis* (11.6% of the total) and 544 genes in *C. teleta* (1.84% of the total) showed large methylation changes, with 51 and 1,809 genes being hyper- and hypomethylated, respectively, in *O. fusiformis* between adult and larval stages, and 125 and 419 genes being hyper- and hypomethylated in *C. teleta* at similar time points (Fig. S8b, d). Transcriptionally, genes with decreased GbM in the adult behave like the other methylated genes whose methylation status remains stable. In contrast, unmethylated genes in the adult stage show a dynamic expression profile during the life cycle of *O. fusiformis*, with a sustained increase in average transcriptional levels from gastrulation to the juvenile and adult stages, thus indicating that GbM dynamics are not linked to transcriptional changes (Fig. S9a), and that genes expressed later in development do not require GbM. In *C. teleta*, however, all genes increase their median transcriptional levels as the life cycle progresses, irrespectively of GbM (Fig. S10a). In both *O. fusiformis* and *C. teleta*, the genes with variable GbM levels in the adults are related to diverse biological processes, from various metabolic processes in *O. fusiformis* (Fig. S9b, c) to DNA recombination and regulation in *C. teleta* (Fig. S10b, c). Therefore, changes in GbM during the annelid life cycle––particularly in the adult stage––are unlikely to result from the activity of specific developmental programmes and present a poor link with transcriptional changes.

Global and gene-level methylation levels decrease during annelid development, yet the loss of methylation might occur preferentially in some regions, e.g. *cis*-regulatory elements [15–17, 23]. To assess this, we identified differentially methylated regions (DMRs) between the three developmental time points, and consistent with the global loss of DNA methylation, the vast majority of DMRs (86.3% and 73.4%) become hypomethylated in *O. fusiformis* and *C. teleta* adult stages, respectively (Fig. 2f). As expected by the extent of 5mC loss between adult and larval stages, the overall number of DMRs is more pronounced in *O. fusiformis* than in *C. teleta* (Fig. 2f). The majority of DMRs (83%) in *O. fusiformis* occur within non-coding (intergenic or intronic) and potentially non-regulatory regions, as they do not overlap with open chromatin as determined by existing ATAC-seq datasets (11.5% DMRs overlap distal ATAC peaks; Fig. 2g). In most of these cases (89.4%), the DMRs contain a TE. In *C. teleta*, hypomethylated DMRs in the adult are also most abundant in non-regulatory, intergenic regions (30.2%), followed closely by promoter regions and transcribed units (exons and introns) (Fig. 2h). As in *O. fusiformis*, the majority of DMRs in intergenic (76.3%), intronic (68 .9%) and promoter (60.7%) regions in *C. teleta* include a TE (Fig. 2h). Notably, the temporal expression dynamics of the genes with a hypermethylated DMR in their promoters in the adult stage was not overtly different from those with a hypomethylated promoter DMR in any of the comparisons and annelid species (Fig. S11a–d). Likewise, these genes participate in various biological processes in both annelids (Fig. S12). Therefore, these findings reinforce the observation based on GbM variation (Fig. S9, S10) that changes in methylation status probably have context-specific or no direct causal effects on transcription in annelids, not following a straightforward link with developmental transcriptional repression or activation.

### DNA methylation is associated with intronic transposable elements in O. fusiformis

DNA methylation regulates the activity of transposable elements (TEs) in various animal lineages [50–53]. Because methylation levels decrease at adult stages in annelids, we tested whether this could be linked to TE reactivation in these organisms. Consistently, in *O. fusiformis* and *C. teleta*, the greatest changes in TE expression occur between adult and larval stages. However, thousands of TEs are up or downregulated (Fig. S13a, f) regardless of their position and family, and there is not a general trend towards global TE upregulation (Fig. S13b, c, e, g, h, i), which suggest a weak relationship between broad methylation levels and TE activity in these annelids (Fig. S14), a link probably restricted to some specific contexts.

To discriminate whether 5mC might specifically target TEs in these annelids, we analysed GbM levels of genes with and without TEs in their introns. In *O. fusiformis* and *C. teleta*, GbM levels increase with the number of introns in an open reading frame (Fig. S15a, c). Likewise, GbM levels in *O. fusiformis* increase as the number of TE-containing introns rises (Fig. S15b). In *C. teleta*, however, this correlation is not obvious. While *C. teleta* genes with 1–5 TE-containing introns have higher GbM than those without introns, the GbM levels of genes with >7 TE-containing introns are low (Fig. S15d). Notably, the presence of a TE-containing intron rapidly raises GbM levels compared to transcriptional units with the same intron number but no TEs in *O. fusiformis*, which again only applies to genes with fewer introns in *C. teleta* (Fig. 2i; Fig. S15e, f). Therefore, these data indicate that GbM levels are linked to the presence/absence of TEs in *O. fusiformis*, suggesting that they might be specifically targeted in this annelid species. However, this is probably not a widespread mechanism across invertebrates, as previously proposed [52].

### Changes in transcriptional dynamics explain interspecies differences in DNA methylation

While the link between intronic TEs and GbM is restricted to *O. fusiformis*, the association between GbM and transcriptional dynamics occurs in this annelid and *C. teleta*. GbM has been proposed to conserve transcriptional dynamics of highly expressed genes, mainly with housekeeping functions, in a few invertebrate lineages [12]. To explore this in annelids, we compared the transcriptional dynamics and GbM levels of 4,460 one-to-one orthologues between *O. fusiformis* and *C. teleta*. Most genes show conserved GbM status between the two species at gastrula (stage with highest 5mC levels) and adult (stage with lowest 5mC levels) stages (Fig. 3a). However, 11.5% of the orthologues are hypermethylated in *O. fusiformis* compared to *C. teleta* in adults, but only 1.2% are hypermethylated in *C. teleta* in the same stage, in agreement with the generally lower global GbM levels in this species (Fig. 3a). Interspecies differences in GbM affect broad functional gene categories, as identified through the enrichment of GO categories. Terms related to development, neurogenesis and core cellular processes are overrepresented in the hypermethylated genes in the adult *O. fusiformis* (Fig. S16a). In contrast, overrepresented GOs are related to immunity, RNA biology, and metabolism in *C. teleta* (Fig. S16b). Notably, and as previously observed [29], pairs of one-to-one orthologs tend to diverge in their transcriptional dynamics as the life cycle progresses in these two annelids (Fig. 3b). In contrast, the correlation of GbM levels between these gene pairs increases from gastrula to adult stages (Fig. 3b), as expected if GbM methylation is more frequently retained in stable, housekeeping genes that might show less interspecific transcriptional differences as development progresses. Indeed, the differences in GbM between *O. fusiformis* and *C. teleta* are partially attributed to interspecific differences in expression levels (i.e., hypermethylated genes in *O. fusiformis* are more highly expressed than their orthologs in *C. teleta*, and vice versa) (Fig. 3c), and transcriptional stability (i.e., hypermethylated genes in *O. fusiformis* are less dynamically expressed during development than their orthologs in *C. teleta*, and vice versa) (Fig. 3d). Therefore, the varying levels of global and GbM found in annelids are at least partially the result of species- and gene-specific transcriptional dynamics, reinforcing the connection between this base modification and gene expression in these organisms, although GbM might be a downstream consequence of gene transcription rather than a developmental regulatory mechanism of particular gene regulatory programmes in these species.

**Figure 3.**
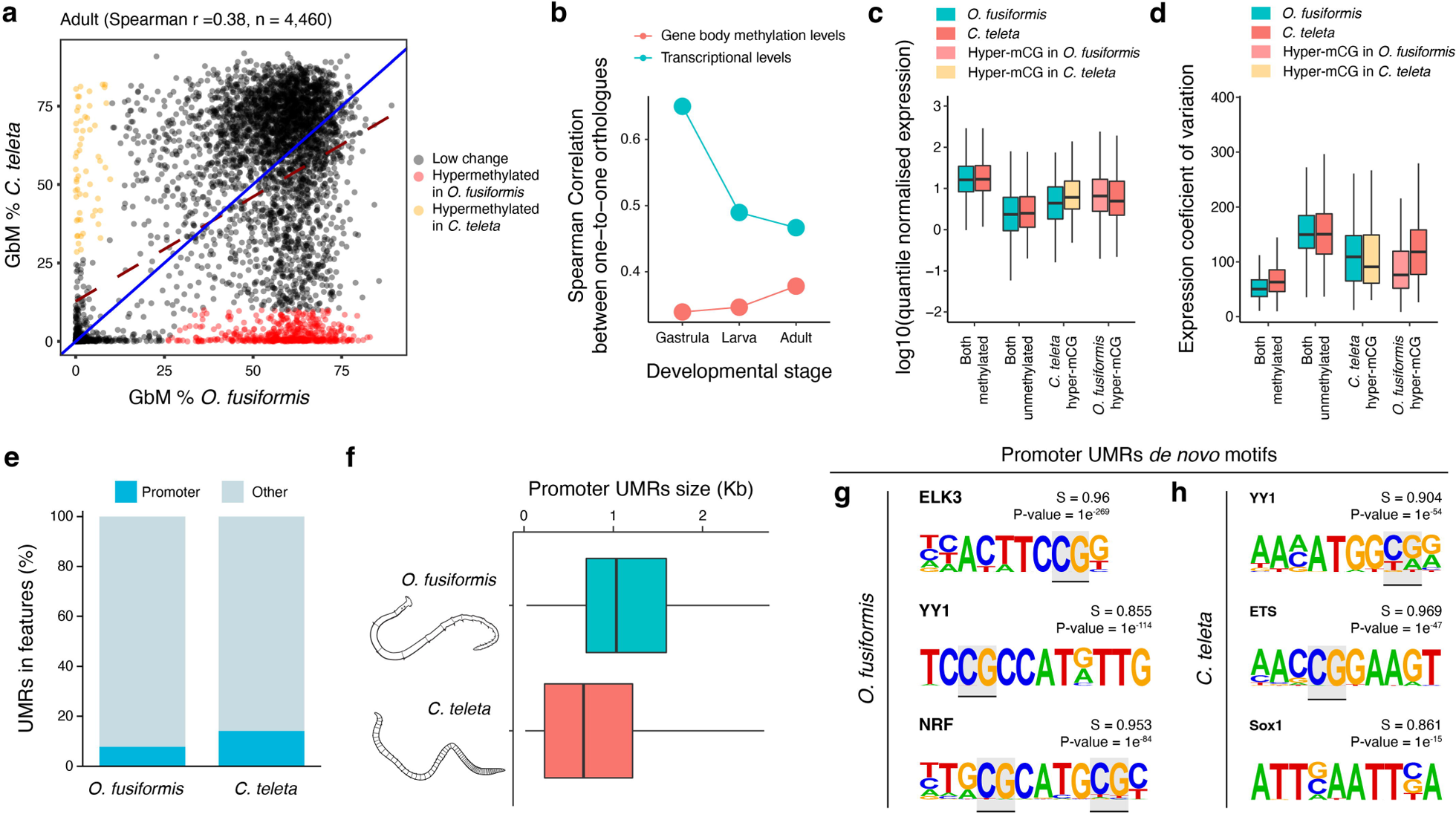
Interspecific differences in DNA methylation dynamics. (**a**) Scatter plots correlating gene body methylation (GbM) levels in one-to-one orthologous genes between *O. fusiformis* and *C. teleta* at the gastrula (left) and adult (right) stages. Hypermethylated genes in *O. fusiformis* compared to *C. teleta* are in red, and hypermethylated genes in *C. teleta* are in yellow. (**b**) Correlation in gene body methylation and transcriptional levels between *O. fusiformis* and *C. teleta* at the three sampled stages. While the correlation in transcriptional levels in one-to-one orthologues decreases as the life cycle progresses, global GbM patterns become more similar with age between the two annelids. (**c**) Box plots of the distribution of expression values of pairs of one-to-one orthologues that are methylated (≥20% mCG) and unmethylated (<20% mCG) in both annelids, hypermethylated in *O. fusiformis* and hypermethylated in *C. teleta* as per panel (**a**). Consistent with the positive correlation between gene body methylation and expression levels, hypermethylated genes in either of these annelids correlates with higher expression values on that species compared to the other one. (**d**) Box plots of the distribution of expression values of pairs of one-to-one orthologues that are methylated and unmethylated in both annelids, hypermethylated in *O. fusiformis* and hypermethylated in *C. teleta*. Consistent with the positive correlation between gene body methylation and expression stability, hypermethylated genes in either annelid correlate with lower transcriptional variation on that species compared to the other one. (**e**) Bar plots showing the proportion of unmethylated regions (UMRs) in promoters and other genomic regions in *O. fusiformis* and *C. teleta*. Only a small fraction of UMRs colocalise with promoters. (**f**) Box plots showing the size distribution of promoter UMRs in the two studied annelids. Consistent with their different genome sizes, *O. fusiformis* has larger promoter UMRs than *C. teleta*. (**g**, **h**) Transcription factor DNA binding motifs enriched in promoter UMRs in *O. fusiformis* (**g**, left) and *C. teleta* (**h**, right). Promoter UMRs are enriched in methylation-sensitive DNA binding motifs (potentially methylated CGs are highlighted in gray).

Depletion of 5mC in upstream promoter regions is a common feature in many organisms that influences the binding affinity of methylation-sensitive transcriptional regulators [7, 54], which might ultimately explain the inter-specific differences in transcriptional dynamics and GbM between species. As such, most unmethylated regions (UMRs) in, for example, the hypermethylated genomes of sponges and vertebrates occur in promoters [7, 9]. In *O. fusiformis* and *C. teleta*, however, only 7.71% and 14.08% of the UMRs correspond with promoter regions (Fig. 3e), respectively (Fig. 1g). In agreement with the different genome sizes between these two annelids (Fig. 1b), the size of promoter UMRs is larger in *O. fusiformis* than in *C. teleta* (Fig. 3f). Further, in both species, these regions are enriched in transcription factor DNA binding motifs robustly annotated to 5mC-sensitive transcriptional regulators, such as ELK3, YY1, NRF and ETS (Fig. 3g, h), as in other species [54]. Therefore, transcription factor methyl-sensitivity shapes the regulatory information found in unmethylated promoters, which together with the different dynamics of genome regulation during their embryogenesis [29], might contribute to the gene- and species-specific differences in DNA methylation between these two annelids.

### A gradual, age-dependent erosion of the post-embryonic methylome in *C. teleta*

The global depletion of 5mC from embryonic to adult stages in *O. fusiformis* and *C. teleta* suggests that these organisms experience a developmental erosion of their methylome. However, this effect could extend post-embryonically in an ageing-associated manner, as it is well established for vertebrates [55–57]. To test this, we generated replicated low-coverage Nanopore methylomes at four ages in *C. teleta*, as its entire life cycle, from egg to senescent adult, can easily be followed under laboratory conditions and lasts around six months. This annelid has an indirect development involving a lecithotrophic larva that, in its latest stages before metamorphosis, already shows adult-like characters, such as a segmented trunk (Fig. 4a) [47]. Thus, metamorphosis is minimal and results in a juvenile worm that burrows and feeds in mud (Fig. 4b). After two months with *ad libitum* food and at 15°C, these juveniles mature sexually and reach their reproductive peak at around three months post-metamorphosis (Fig. 4c). Adults of *C. teleta* are iteroparous and reproduce several times during their lifetime, experiencing a progressive age-dependent anatomical decay (Fig. 4d). Consistent with the data from EM-seq (Fig. 2b), global 5mC levels gradually decrease from the larva to juvenile stages in *C. teleta*. However, the loss of methylation is more acute as the worm undergoes several reproductive cycles and enters senescence (Fig. 4e). Therefore, post-embryonic age-dependent demethylation occurs in *C. teleta*, and perhaps as well in other annelids like *P. dumerilii* [8], suggesting that the link between ageing and epigenetic erosion might exhibit broad commonalities between distantly phylogenetically related animals.

**Figure 4.**
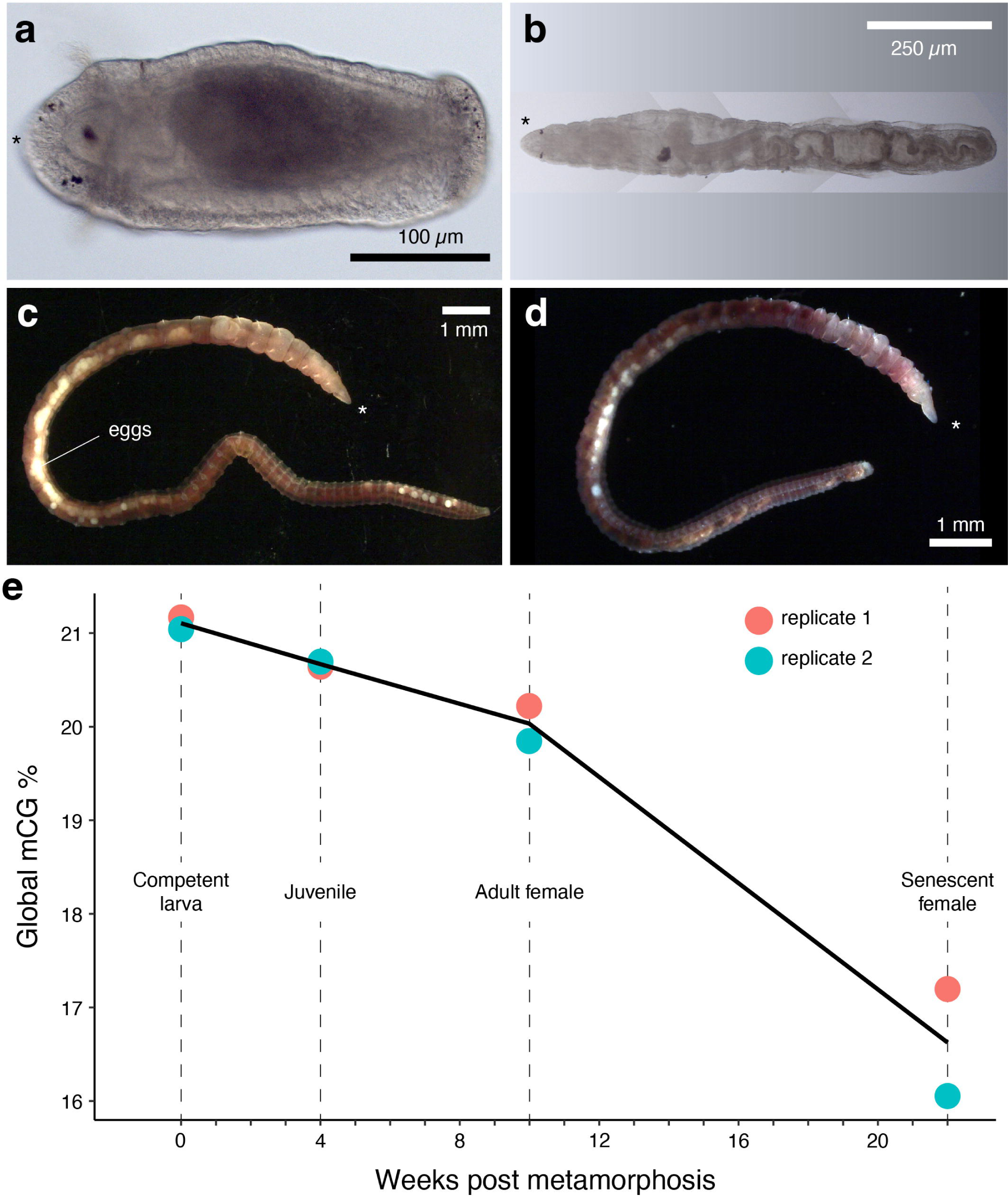
A gradual global demethylation in adult *C. teleta* with ageing. (**a,b,c,d**) Photographs of the four sampled stages. (**e**) Line plot of global methylation levels obtained from multiplexed Nanopore libraries sequenced at shallow coverage from late larval stages until ageing adults. Asterisk in **a**–**d** denote the anterior end.

### Cytidine analogues disrupt embryogenesis in O. fusiformis

As an approach to functionally test the role of 5mC during annelid development, we treated fertilised oocytes of *O. fusiformis* until larval development with zebularine, 5-azacytidine, and decitabine, three cytidine analogues that impair DNA methylation by incorporating into DNA and blocking DNMTs (Fig. 5a) [58–60]. Unlike in the annelid *P. dumerilii* [8], decitabine was toxic at all tested concentrations, blocking the division of the zygote. Thus, we did not use it in further analyses. With zebularine treatment, most treated embryos (82.87%) developed into abnormal larvae in a dose-dependent manner (Fig. 5b; Fig. S17a; Table S3). As the control larvae, zebularine-treated larvae are bilaterally symmetrical and have a U-shaped gut and a blastocoel. Yet, treated larvae are smaller and less elongated than controls and have general differentiation problems, lacking a well-developed locomotory ciliated band, a normal number of posterior defensive chaetae, a sensory apical organ, and a prominent stomach (Fig. 5b; Fig. S17a). Treatment with 5-azacytidine either killed the embryo (25.23%) or resulted in abnormal cleavage and gastrulation failure (74.77%), with the embryos becoming a disorganised mass with some ciliated cells (Fig. S17a), indicating that 5-azacytidine, as decitabine, is probably toxic in this annelid. Only zebularine treatment causes a modest, yet significant, depletion (*p*-value = 0.0035; two-sided t-test) of global 5mC levels during *O. fusiformis* embryogenesis (Fig. 5c). Therefore, disruption of DNA methylation levels with zebularine leads to defective embryogenesis in the annelid *O. fusiformis,* although other cytidine analogues cause severe phenotypes without a clear impact on 5mC.

**Figure 5.**
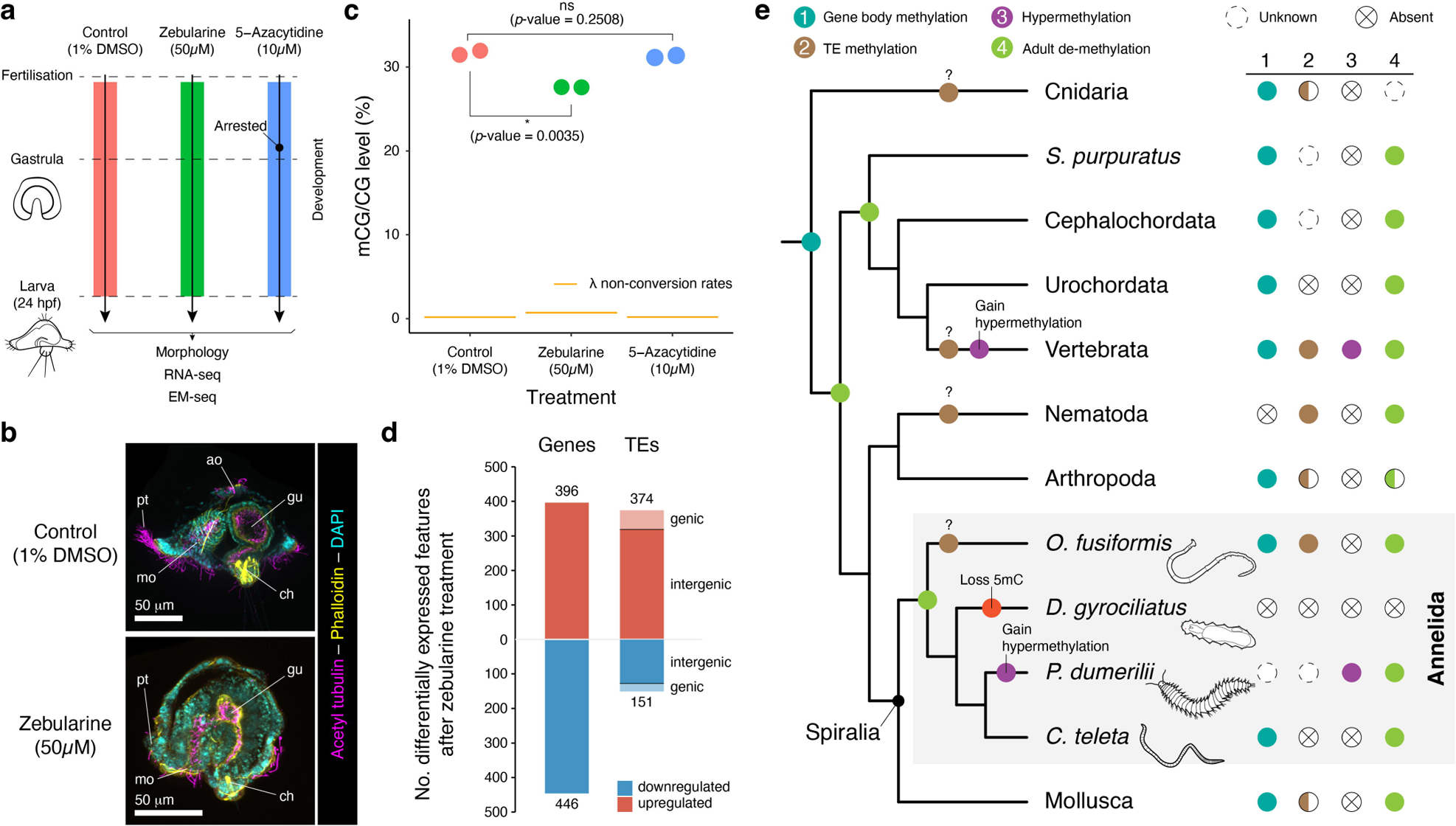
DNA methylation is essential for normal embryogenesis in *O. fusiformis*. (**a**) Schematic representation of the experimental design. Independent batches of embryos were treated after fertilisation with either 1% DMSO (control) or 50 µM zebularine and 10 µM 5-azacytidine for 24 hours until the larval stage when phenotypes were scored. (**b**) Z-projections of confocal stacks of zebularine-treated and DMSO-control embryos fixed at the early larval stage and stained for acetylated tubulin (magenta), actin (yellow) and nuclei (cyan). Zebularine treatment impairs annelid embryogenesis as treated larvae fail to undergo normal organogenesis in *O. fusiformis*. (**c**) Zebularine but not 5-azacytidine treatment significantly decreases global methylation levels in *O. fusiformis* as measured with shallow coverage EM-seq. (**d**) Bar plots depicting the number of differentially expressed genes and transposable elements (TEs) after zebularine treatment. Reduction of normal methylation levels reactivates TE expression, mainly in intergenic regions. (**e**) The evolution of DNA methylation in bilaterally symmetrical animals and Spiralia. Gene body methylation is likely the ancestral stage in animals and the presence of adult de-methylation in deuterostomes and annelids suggests that this might be an ancestral bilaterian feature. TE methylation might have evolved multiple times independently, similarly to the levels of DNA methylation, with multiple independent transitions to a hypermethylation and unmethylation state in different animal lineages.

As expected from the treated phenotypes, zebularine impairs normal gene expression in *O. fusiformis*. Replicated transcriptomes of treated and control larvae (Fig. S17b, c) revealed 396 upregulated and 446 downregulated genes after zebularine treatment (Fig. 5d). Upregulated genes exhibit higher average 5mC levels than downregulated genes (Fig. S17d), and accordingly, they display more stable transcriptional dynamics during embryogenesis (Fig. S17e). While the upregulated genes are enriched in GO terms associated with morphogenesis and stimuli response, the downregulated genes are related to stress and ion homeostasis (Fig. S17f, g), which might explain the smaller volume of the internal fluid-filled blastocoel. Zebularine treatment, however, largely upregulates TE transcription in *O. fusiformis* (Fig. 5d). Notably, most of the dysregulated TEs (85.29% of the upregulated and 84.11% of the downregulated) occur in intergenic regions (Fig. 5d), suggesting that their change in expression is unrelated to differences in gene transcription. In addition, LTRs are the largest upregulated TE class after zebularine treatment (Fig. S17h). However, these are not the most abundant in the genome of *O. fusiformis* [29], which might reflect some degree of specificity in the regulatory impact of 5mC on TE activity in this annelid species. Our findings thus functionally support that DNA methylation is involved in gene and TE regulation during annelid embryogenesis. This coupled with previous findings during posterior regeneration in the annelid *P. dumerilii* [8] indicate that normal levels of 5mC are important for successfully deploying genetic programmes during the annelid life cycle.

## Discussion

This study uses genome-wide, base-resolution profiling to characterise the landscape, dynamics, and function of DNA methylation during the life cycles of three annelids with distinct genomic features and phylogenetic positions, allowing us to reconstruct ancestral and derived traits for this base modification in a major invertebrate clade (Fig. 5e). All possible patterns of 5mC global methylation occur in Annelida, from the potentially hypermethylated genomes of some species, like *P. dumerilii* [8, 36], to the more typical invertebrate low-to-mid levels of *O. fusiformis* and *C. teleta*, and the loss of methylation in *D. gyrociliatus*. Given the DNA methylation levels of molluscs and other invertebrate groups [1, 2, 37], a hypermethylation state is likely a secondary modification in Annelida restricted, so far, to a single clade (Nereididae) (Fig. 5e), which will require confirmation with genome-wide techniques. In *D. gyrociliatus*, the secondary loss of 5mC (Fig. 5e) correlates with a modified DNA methylation toolkit, particularly with the absence of DNMT3 and a divergent N-terminus for DNMT1, which lacks the autoinhibitory zinc finger CXXC domain that recognises unmethylated CGs [61, 62]. Nonetheless, this DNMT1 retains potential catalytic activity in its methyltransferase domain and an N-terminus Replication Foci Domain (RFD), which drives the localisation of DNMT1 to the replication forks. This is reminiscent of the beetle *Tribolium castaneum*, whose genome also encodes a DNMT1 but lacks DNMT3 and similarly shows the absence of 5mC [63]. Still, in *T. castaneum*, DNMT1 knockdown impairs embryogenesis, suggesting that this gene plays important roles beyond 5mC deposition in invertebrates and that DNMT1 loss might have pleiotropic effects in some lineages beyond controlling gene expression and 5mC maintenance [64].

As the life cycle progresses, the DNA methylation landscape erodes in *O. fusiformis* and *C. teleta*, with global and CpG level methylation levels falling along development and the corelation with transcription weakening. This is in line with decreased methylation levels between larval and juvenile stages in *P. dumerilii* detected by luminometric methylation assay [8]. 5mC erosion is consistent with inverse temporal patterns of gene expression between DNA methylation writers (DNMT1 and DNMT3), prevalent in early development, and erasers (TET), which upregulate after gastrulation and are highly and broadly expressed in later life stages. Similar adult demethylation and DNMT–TET transcriptional dynamics occur in vertebrates and related deuterostome invertebrates, such as amphioxus and sea urchins [16, 17, 23], suggesting that global demethylation as development progresses is a conserved epigenomic trait, at least for bilaterians (Fig. 5e). In vertebrates, this demethylation occurs in regulatory regions of developmental genes [16, 17, 23]. In deuterostome invertebrates, some of this demethylation occurs in intra-genic regulatory regions marked by ATAC-seq, but it is also widespread across the genome [23]. Although some regions that become hypomethylated during development overlap with potential *cis*-regulatory elements (ATAC-seq peaks) in annelids, this link appears weaker than in deuterostomes. Indeed, most hypomethylated DMRs in adult stages occur in intergenic regions and intronic regions not overlapping ATAC peaks in *O. fusiformis* and *C. teleta*, and affect genes involved in diverse biological processes not restricted to developmental functions. This indicates that adult demethylation is probably a global dynamic rather than a tightly regulated process linked to the unfolding of specific developmental programmes in these animals. In support of this scenario, TET proteins in annelids and other spiralians, but not deuterostomes, lack the N-terminus zinc finger CXXC domain (Fig. 1c) [7, 8, 23, 65, 66], which would confer DNA binding specificity to this DNA methylation eraser [67]. Age-dependent changes in DNA methylation status at specific CpGs are good predictors of biological age in mammals [68, 69], and global, genome-wide hypomethylation is a landmark of senescence in many vertebrate cell types [70]. Notably, the gradual, age-dependent DNA methylation erosion observed in annelids accelerates with ageing, at least in *C. teleta*. Therefore, comparable manifestations of senescence occur in distantly related animals, opening the possibility that chronological “epigenetic clocks” [71] might exist more broadly than previously recognised in animals. More importantly, future studies comparing methylation across wild individuals should consider age as a crucial variable explaining global and GbM levels in invertebrates.

As in most invertebrates with mosaic patterns of DNA methylation [1, 2, 6], 5mC accumulates in gene bodies, correlating positively with transcriptional activity and stability in annelids. Although the functional role of GbM is poorly understood, one of the main theories posits that it prevents spurious transcriptional initiation from otherwise highly transcribed regions [20]. The fact that intronic TEs influence GbM, at least in *O. fusiformis*, could support this hypothesis, as potentially, the TEs that land in these methylated regions are less likely to be detrimental and more likely to be kept, or DNA methylation is actively targeting these genes to avoid TE transcription [20]. However, *C. teleta* does not show an association between intronic TEs and GbM, indicating that the link between these features is probably evolutionary plastic, as previously noted in cnidarians [52]. GbM might also be important to keep high and robust transcriptional levels of the methylated genes in invertebrates [12]. This correlation with transcription stability is present in *O. fusiformis* and *C. teleta*, and the orthologues that change GbM levels across species also show divergent transcriptional levels and dynamics (loss of methylation linked to higher coefficient of variation). However, the zebularine treatment decreases global methylation in *O. fusiformis*, yet does not result in general repression of genes with GbM. On the contrary, genes upregulated upon zebularine treatment show slightly higher methylation levels than those downregulated. This starkly contrasts what was observed after DNMT1 knockdown in the wasp *Nasonia vitripennis*, where the loss of GbM was associated with transcriptional downregulation, whereas unmethylated genes were upregulated [72]. This indicates that the causal link between GbM and transcription is still poorly understood in invertebrates and will require precise gene disruption strategies to disentangle the various proposed functions.

5mC TE methylation has been described in diverse animal taxa, from cnidarians to vertebrates and nematodes [11, 52, 53, 73] (Fig. 5e) and proposed to be a mechanism to suppress TE activity, particularly in lineages with high transposon burden [74]. However, the relationship between DNA methylation and TEs appears species-specific in most invertebrates. For instance, in molluscs, DNA methylation outside of gene bodies occurs preferentially in some young TE classes in the bivalve *C. gigas*, yet the TE-rich genomes of cephalopods do not show strong TE methylation [39, 41]. Likewise, among all studied annelids, the association between DNA methylation and TEs, regardless of the class, is more pronounced and perhaps unique to *O. fusiformis* (Fig. 5e) [42]. Indeed, TE load does not correlate well with global 5mC levels and TE methylation in annelids: the deep-sea species *Paraescarpia echinospica* has a higher TE burden than *O. fusiformis* but does not show TE methylation [29, 42, 75]. Alternative pathways to control TE activity, such as PIWI-interacting RNAs (piRNAs) [76, 77], are more broadly conserved in annelids and molluscs [78, 79]. In addition, although the mechanism of how DNMTs target gene bodies is quite well understood (the PWWP domain in DNMT3 orthologues binds to histone lysine 36 trimethylated residues typical of gene bodies) [80], how invertebrate DNMTs might target TEs remains unresolved. In tetrapods, for instance, DNMTs are targeted to TEs guided by KRAB zinc fingers, a fast-evolving transcription factor family that recruits various chromatin modification enzymes [81, 82]. However, vertebrate genomes are mostly hypermethylated by default; thus, targeting both old and new TEs is easier to explain. Species like *O. fusiformis*, in which there is some degree of intergenic TE methylation, will be the best suited to reveal how TEs are targeted in genomes with mosaic methylation patterns. Nonetheless, given the current sampling among invertebrates and the relative scarcity of strong intergenic TE methylation in invertebrates, the convergent evolution of this mechanism across invertebrates and vertebrates from an ancestral animal pattern mostly restricted to gene bodies is the most parsimonious scenario (Fig. 5e).

Successful posterior adult regeneration and normal embryogenesis are impaired when using cytidine analogues in the annelid *P. dumerilii* and the bivalve *C. gigas*, respectively [8, 40]. In the annelid *O. fusiformis*, decreasing global DNA methylation levels with the cytidine analogue zebularine also affects normal organogenesis, resulting in well-patterned larvae with immature organs. Notably, the phenotype described in *C. gigas* with 5-azacytidine–– exogastrulation and development into disorganised ciliated cells [40]––resembles the outcome observed in *O. fusiformis* after an equivalent treatment, but it is likely a toxic effect because 5–azacytidine treatment does not affect normal 5mC levels. Unlike previous functional works in free-living spiralians, our study explores both the amount of 5mC loss upon treatment and the transcriptional consequences of cytidine analogues during embryogenesis in *O. fusiformis*. Consistent with the TE methylation observed in this annelid, genome-wide hypomethylation during embryogenesis reactivates TEs, particularly LTRs in intergenic regions, but also dysregulates transcription. These significant changes in gene transcription levels do not appear to affect specific gene programmes or genes with determined GbM levels, and result in overall even up- and down-regulation of genes. This is also in accordance with the observed mild developmental phenotype, which affects the general morphology of the larva rather than the formation of specific tissues and organs, thus not suggestive of a role of 5mC in controlling regulatory networks. However, cytidine analogues are known to have toxic off-target effects. Indeed, we observe strong phenotypes in *O. fusiformis* when using 5-Azacytidine without observing global changes in 5mC, which is a cautionary tale when performing this kind of treatment in invertebrates. Further, the reduction in global methylation levels when using zebularine is modest (from 32 to 28%), implying a lot of cell-to-cell variability in methylation loss, hampering the resolution of how this loss might influence transcription.

## Conclusions

Our comprehensive, base-resolution characterisation of the dynamic DNA methylation landscape during the life cycle of three annelids expands our knowledge of the evolution and roles of this pervasive base modification in one of the largest invertebrate clades. While a mosaic pattern of DNA methylation concentrated in active gene bodies is ancestral in annelids, this landscape has diverged in some lineages, with cases of potentially hypermethylated (Nereididae) and unmethylated (*D. gyrociliatus*) genomes. Likewise, methylation of transposable elements occurs in some lineages (*O. fusiformis*) but not others. However, the erosion of the DNA methylation landscape as the life cycle progresses is conserved in Annelida, and since it has also been observed in deuterostome invertebrates and insects, we suggest that developmental epigenetic erosion is an ancestral feature of bilaterians. Postembryonic methylation loss is associated with ageing in the annelid *C. teleta*, implying that epigenetic clocks can be obtained in non-vertebrates. In the future, developmental genome-wide, base-resolution methylomes of more species, particularly those with divergent patterns, combined with direct manipulation of methylation patterns with gene disruption approaches, will provide a more complete view of the functional implications of DNA methylation in annelids and invertebrate genomes.

## Methods

### Animal husbandry and embryo collection

Sexually mature *Owenia fusiformis* (Delle Chiaje, 1844) adults were acquired from the coasts near the Station Biologique de Roscoff (France) during the reproductive season and cultured in artificial seawater (ASW) at 15°C as previously described [83]. Following *in vitro* fertilisation [83], embryos were incubated and left to develop at 19°C in filtered ASW until the desired embryonic and larval stages. *Capitella teleta* (Blake, Grassle & Eckelbarger, 2009) adults were cultured at 19°C in ASW with embryonic and larval stages collected based on previously established protocols [47]. *Dimorphilus gyrociliatus* (O. Schmidt, 1857) were also cultured at 19°C in a 4:5 ratio of ASW and freshwater, respectively [45].

### Orthology assignment and domain architecture analyses

DNMT, MBD, and TET sequences were mined from genomic and transcriptomic resources for *O. fusiformis*, *C. teleta* and *D. gyrociliatus* [29, 45] and aligned against other representative sequences with MAFFT v7.505 in L-INS-I strategy [84]. Resulting alignments were manually trimmed in Jalview v2.11.2.5 [85] according to the main domain boundaries of each gene family (PF00145 for DNMT, PF01429 for MBD, and PF12851 for TET), followed by removal of poorly aligned regions with TrimAl v1.4.rev15 in automated mode [86]. Maximum likelihood trees were then constructed with IQ-TREE v2.2.0.3 [87] using 1,000 ultrafast bootstraps and the “-m TEST” option to identify the best amino acid substitution model for each gene family. Bayesian reconstruction was performed with MrBayes v.3.2.7a [88] using the LG+Gamma model (for DNMTs and TETs) and GTR (for MBDs) until convergence. All trees were visualised and edited in FigTree v.1.4.4 (https://github.com/rambaut/figtree/). The protein domain composition of each gene was characterised with InterProScan5 [89] and constructed in IBS 1.0 [90].

### Gene expression profile of DNA methylation-related genes

Expression dynamics of DNMT1, UHRF1, DNMT3, and TET genes were retrieved from publicly available temporal RNA-seq series for *O. fusiformis*, *C. teleta* and *D. gyrociliatus* [29, 45]. Additionally, replicated RNA-seq libraries were constructed for the adult stages of *O. fusiformis* and *C. teleta*. Total RNA was extracted with the RNA Miniprep Kit (New England Biolabs, #T2010) according to the manufacturer’s instructions and used for standard strand-specific Illumina library prep, which was sequenced in a NovaSeq 6000 platform in pair-end 150 bases mode. Sequencing adaptors were trimmed with fastp v0.20.1 [91] and pseudo-aligned to gene models with Kallisto v0.46.2 [92]. DNA methylation-related gene expression profiles of *O. fusiformis* and *C. teleta* were plotted in R using ggplot2 v3.4.0 [93] based on quantile normalised transcripts per million (TPM) to account for technical and biological variations between libraries. Normalised counts were used to plot these gene expression profiles for *D. gyrociliatus*.

### Whole-mount in situ hybridisation

DNMT1, DNMT3, MBD1/2/3 and TET genes were amplified from cDNA containing various developmental stages of *O. fusiformis* through two successive rounds of nested PCR that added a T7 promoter at the 3’ end of the amplicon. The resulting DNA templates were *in vitro* transcribed to obtain DIG-labelled riboprobes using T7 enzyme (Ambion MEGAscript kit, #AM1334) and stored at −20°C in hybridisation buffer at a total concentration of 50 ng/μl. Colourimetric whole-mount *in situ* hybridisation was performed following established protocols [44]. Representative samples were imaged with a Leica DMRA2 upright microscope and Infinity5 camera (Lumenera) using differential interference contrast (DIC) options. Resulting whole images were adjusted for brightness, contrast and colour balance in Adobe Photoshop and assembled into a final figure panel in Adobe Illustrator.

### DNA methylation sequencing

Whole genome bisulphite sequencing (WGBS) libraries were constructed for *O. fusiformis* (gastrula, larva and adult) and *D. gyrociliatus* (adult) samples with single replicates each (Table S1). Bisulphite conversion was performed using the EZ DNA Methylation Gold Kit (Zymo Research), followed by size selection and library amplification. Developmental time points (gastrula, larva and adult) of *C. teleta* were assayed by Enzymatic-Methyl seq (EM-seq) (NEB #E7120) according to the manufacturer-provided protocol. For EM-seq negative and positive controls, spikes of unmethylated lambda phage DNA and methylated pUC19 DNA were added before enzymatic treatment, whereas only lambda phage DNA was used in WGBS. All the resulting libraries were sequenced in 150-bases paired-end mode on an Illumina platform. Adaptors were trimmed with fastp v0.20.0 [91], aligned to their respective reference genomes, deduplicated and methylation called for all cytosine contexts (CpG, CHG, and CHH) using Bismark v0.22.3 [94] with default options. Methylation signals were visualised in IGV v2.8.0 genome browser using bigwig files generated from Bismark bedGraph and coverage files via UCSC *bedGraphToBigWig* function. Visualisations of genomic tracks were plotted with pyGenomeTracks [95]. Bismark text files containing methylated and unmethylated cytosines were converted into CGmap files with CgmapTools v0.1.2 [96] and processed into bsseq objects in R for all further downstream analyses. Gene body methylation values were filtered for minimum mean read coverage of four, and minimal number of five CpGs per gene. Non-conversion rates were calculated based on the methylation levels of spike-ins, mitochondrial genomes and non-CpG contexts.

To characterise the age-dependent global demethylation in *C. teleta*, replicated samples of competent larvae (stage 8), one-month juveniles, gravid females and 22-week-old senescent females were collected. Genomic DNA was extracted with the MagAttract kit (Qiagen) and Nanopore libraries were built with the Rapid barcoding kit (SQK-RBK114.24) following manufacturer’s instructions and sequenced on MinION Mk1C using a MinION R10.4.1 flow cell (FLO-MIN114). The sequencing reads were then base-called and demultiplexed with Guppy (v6.2.1) using the super accuracy model dna_r10.4.1_e8.2_400bps_modbases_5mc_cg_sup.cfg model for methylated CpGs. Global methylation levels were calculated as previously described [97].

### Gene body methylation profiling

Positional heatmaps of mCG/CG levels in protein-coding loci were generated using the bssq R object described above and the *computeMatrix* function of deepTools2 [98], and plotted with *plotHeatmap* and *plotProfile* functions with the “scale-regions” option and a bin size of 100 bases. To correlate gene body methylation with transcription, transcriptomic data in TPM (transcripts per million) for *O. fusiformis* and *C. teleta* was imported and processed in R by ranking the mean TPM for each developmental stage and the coefficient of variation for each gene into deciles using the *quantile* function in R and mean_tpm >= 1 operation to exclude genes with mean TPM expression of less than 1.

Pairwise changes in GbM in both annelids were obtained only for genes with coverage >4 in all samples. We first computed the average difference of GbM across all genes between two stages (e.g. gastrula vs larva) and then identified the genes that deviated twofold from that average to focus on the genes with changes above the global methylation loss. For inter-species comparisons (Fig. 5), we first obtained one-to-one orthologues from an Orthofinder2 run with default diamond parameters and then selected the GbM levels from each orthologue in each species, only accepting orthologues with enough coverage threshold.

### Transposable elements

RepeatModeler2 [99] was used to identify repetitive regions in *D. gyrociliatus*, *C. teleta* and *O. fusiformis*, and the resulting annotations were imported into R as GenomicRanges objects [100], including a filtering step to select for TEs longer than 400bp. We further classified these TEs based on their location in genic or intergenic regions by overlapping them with the gene annotations and identified their methylation status, with values equal to or greater than 20% being defined as methylated. The RepeatModeler2 built-in script to calculate Kimura divergence values was used to estimate the evolutionary divergence of TE families. To investigate the relationship between TE density and GbM levels, we ranked genes based on their number of introns (including the presence or absence of TEs) and corresponding methylation levels using GenomicFeatures v1.48.4 package [100]. TE expression was calculated with TElocal v1.1.1 (https://github.com/mhammell-laboratory/TElocal) and differential expression analysis in a pairwise manner using the DESeq2 v1.36.0 [101] package with the function *lfcShrink*. Differentially expressed TEs were selected based on an adjusted *p*-value ≤ 0.01 and Log2 fold change ± 2.

### Unmethylated regions (UMRs)

Repositories containing genomic information were manually created for *O. fusiformis* and *C. teleta* using the R package BSgenome v1.64.0 [102]. Using BSgenome and methylomes in bsseq object form, UMRs were identified for each species using MethylSeekR v1.36.0 [103] *segmentUMRsLMRs* function with “meth.cutoff = 0.5” and “nCpG.cutoff = 4” parameters. Next, motifs of UMRs less than 5 kilobases overlapping promoter features (2kb upstream and 200bp downstream of the transcription start sites) were analysed using HOMER v 4.11 [104] with *findMotifsGenome.pl* function and motif lengths of 6, 8, 10, and 12.

### Differentially methylated regions (DMRs)

The DSS v2.48.0 R package [105] was used to compute DMRs in a pairwise manner, using the *callDMR* function retaining a minimum number of five CGs for these regions with the minCG=5 and dis.merge=100 parameters. We applied a more stringent delta=0.2 parameter, which specifies a difference in methylation greater than 20%. The resulting DMRs were filtered to ensure a minimum mean coverage of four in these regions, and their genomic distributions were defined using GenomicFeatures v1.48.4 R package [100]. Distal regulatory elements were defined as publicly available consensus ATAC peaks not overlapping promoters (−1000 / + 200 bp from each transcriptional start site) [29]. The developmental expression profiles of the genes with promoter DMRs were plotted based on their log10 transformed stage-specific transcriptomic data. Enrichment of Gene Ontology terms in these genes was calculated with the TopGO R package [106]. All graphs were created with ggplot2 v3.4.0 package [93].

### Inhibitor treatments

Fertilised oocytes of *O. fusiformis* were treated with 50 μM zebularine (Abcam: ab141264) and 10 μM 5-Azacytidine (Abcam: ab142744) using equivalent volumes of dimethyl sulfoxide (DMSO) as a negative control. Treated and control embryos developed for 24 hours at 19°C until the larval stage, when drugs were washed off. Samples collected for RNA-seq and EM-seq protocols were immediately snap-frozen in liquid nitrogen and stored at −80°C before library preparation and sequencing, as detailed above. Samples collected for immunohistochemistry were relaxed in an 8% magnesium chloride solution before fixation in 4% paraformaldehyde for one hour at room temperature (RT). Samples were then washed with 0.1% Tween-20 phosphate-buffered saline (PTw) and stored at 4°C in PTW with 1μM sodium azide.

### Immunohistochemistry

F-actin and antibody staining of control and treated *O. fusiformis* larvae were conducted as described elsewhere [83]. Briefly, samples were permeabilised with PBS + 0.1% Triton X-100LJ+LJ1% bovine serum albumin (PTxLJ+LJBSA) and blocked in PTxLJ+LJ5% normal goat serum (NGS). Samples were incubated with primary antibodies (mouse anti-acetyl-alpha tubulin antibody, clone 6-11B-1 [Millipore-Sigma Aldrich, cat#: MABT868, RRID: AB_2819178] and rabbit anti-FMRF-amide antibody [Immunostar, cat#: 20091, RID: AB_572232]) diluted (1:500) in PTx + NGS overnight at 4°C and washed several times with PTxLJ+LJBSA before overnight incubation with 1:800 diluted Alexa-labelled secondary antibodies (ThermoFisher Scientific), 1:100 Alexa Fluor™ 488 Phalloidin (ThermoFisher Scientific), and 5 µg/ml DAPI in PTx + NGS. Secondary antibodies were washed in PTx + BSA and samples were cleared with 70% glycerol in PBS and stored at 4°C before imaging with a Leica SP5 confocal laser scanning microscope. Resulting z-stack projections were made with Fiji [107], and these images were then edited in Adobe Photoshop and assembled into a final figure panel in Adobe Illustrator.

### RNA-seq profiling and differential expression analyses of control and treated samples

Adapter sequences and poor-quality bases were trimmed with fastp v0.20.1 [91] and aligned to *O. fusiformis* reference genome and annotation [29] (GCA_903813345) using STAR vX [108] and Kallisto v0.46.2 [92], respectively. Resulting bam files were processed with TElocal v1.1.1 (https://github.com/mhammell-laboratory/TElocal) to obtain gene and transposable elements expression counts. Differential expression analyses between zebularine-treated and control samples were performed with the R package DESeq2 v1.36.0 [101] with a significant threshold adjusted to a *p*-value of ≤ 0.01 and a log_2_ fold change of ± 2. A Pearson correlation matrix of the libraries was computed with corrplot v0.92 (https://github.com/taiyun/corrplot), Gene Ontology enrichments were performed with TopGO [106] and all other plots were generated with ggplot2 v3.4.0 [93].

## Abbreviations

5mC: 5-methylcytosine
DMR: differentially methylated region
GbM: gene body methylation
TE: transposable element
UMR: unmethylated region

## Declarations

### Ethics approval and consent to participate

Not applicable

### Consent for publication

Not applicable

### Availability of data and materials

The datasets generated and analysed during the current study are available in Gene Expression Omnibus (GEO). Scripts generated to analyse these datasets are available in a public repository on GitHub (https://github.com/ChemaMD).

### Authors’ contributions

K.G., A. de M., J.M.M.-D., and P.J.H. conceived and designed the study; K.G., Y.L., and J.M.M.-D. collected the samples; K.G., L.S., B.E.D., Y.L., L.X., and J.M.M.-D. extracted and prepared sequencing libraries; K.G., L.S., L.X., A. de M., and J.M.M.-D. did all computational analyses; K.G., and A.M.C.-B. did drug treatments and the characterisation of the resulting phenotypes; F.M.M.-Z. contributed to gene annotation and phylogenetic inferences; K.G., A. de M.. and J.M.M.-D. drafted the manuscript, and all authors critically read and commented on it.

## Acknowledgements

We thank Robert Lowe for early efforts in this project, the staff at the Station Biologique de Roscoff for their help with collection and animal supplies, and the core technical staff at the Department of Biology at Queen Mary University of London and QMUL Genome Centre for their support. This research utilised Queen Mary’s Apocrita HPC facility, supported by QMUL Research-IT (http://doi.org/10.5281/zenodo.438045).

