## Supplementary material for "Annelid methylomes reveal ancestral developmental and ageing-associated epigenetic erosion across Bilateria": Fig. S

**Table of contents**

- **Figure S1–S17.** Supplementary figures 1 to 17.
- **Tables S1–S3.** Supplementary tables 1 to 3.

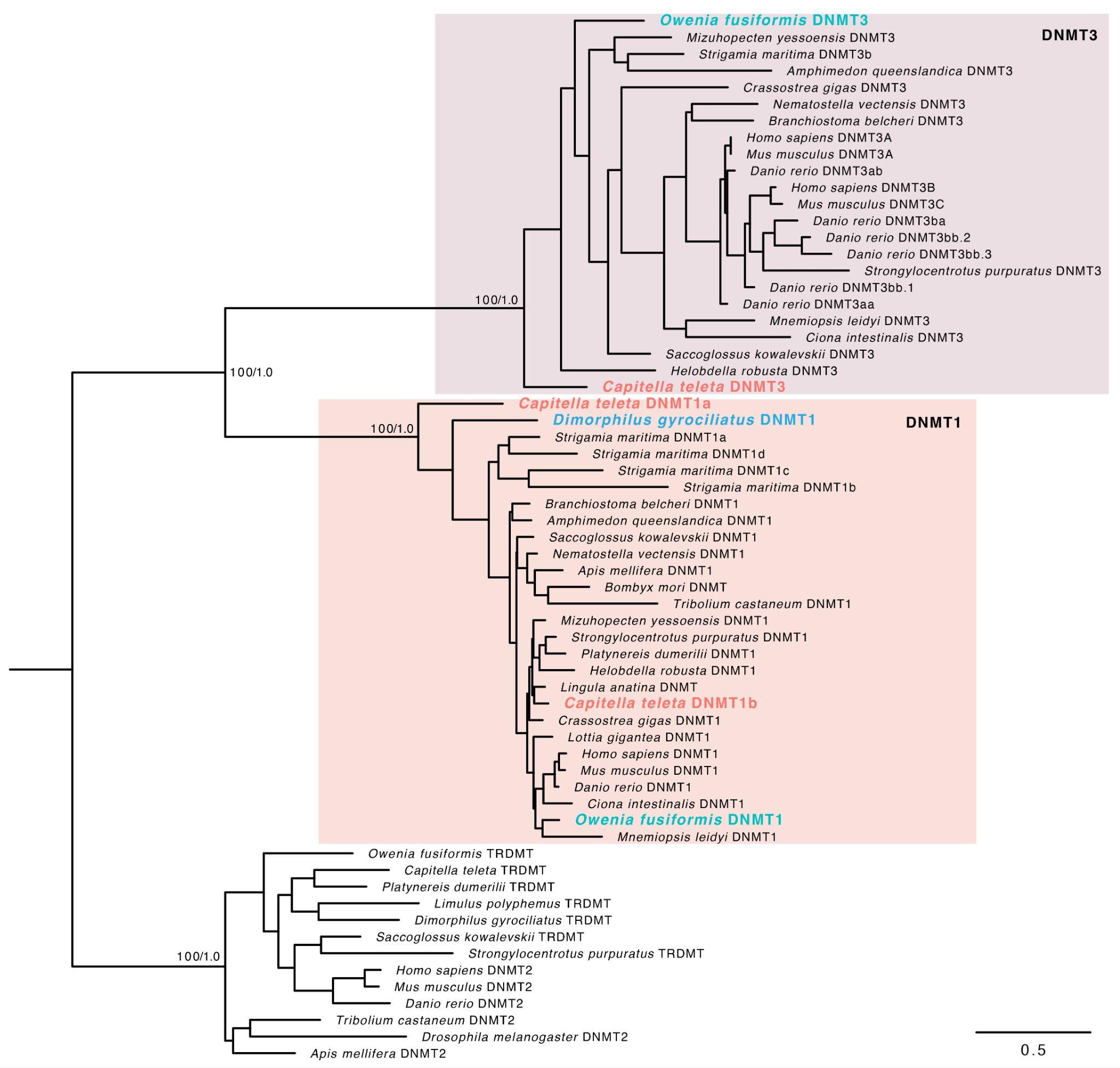

**Figure S1 – Gene orthology assignment of DNMT genes.** Orthology assignment of DNA methyltransferases using the DNMT2 (also known as TRDMT), predominantly methylates tRNAs, as outgroup. The tree topology is based on maximum likelihood reconstruction and node supports indicate both bootstrap values (from 0 to 100) and posterior probabilities (from 0 to 1) at key nodes. Boxes indicate the DNMT1 and DNMT3 sub-families and the scale bar represents the number of amino acids substitutions per site alongside the branches.

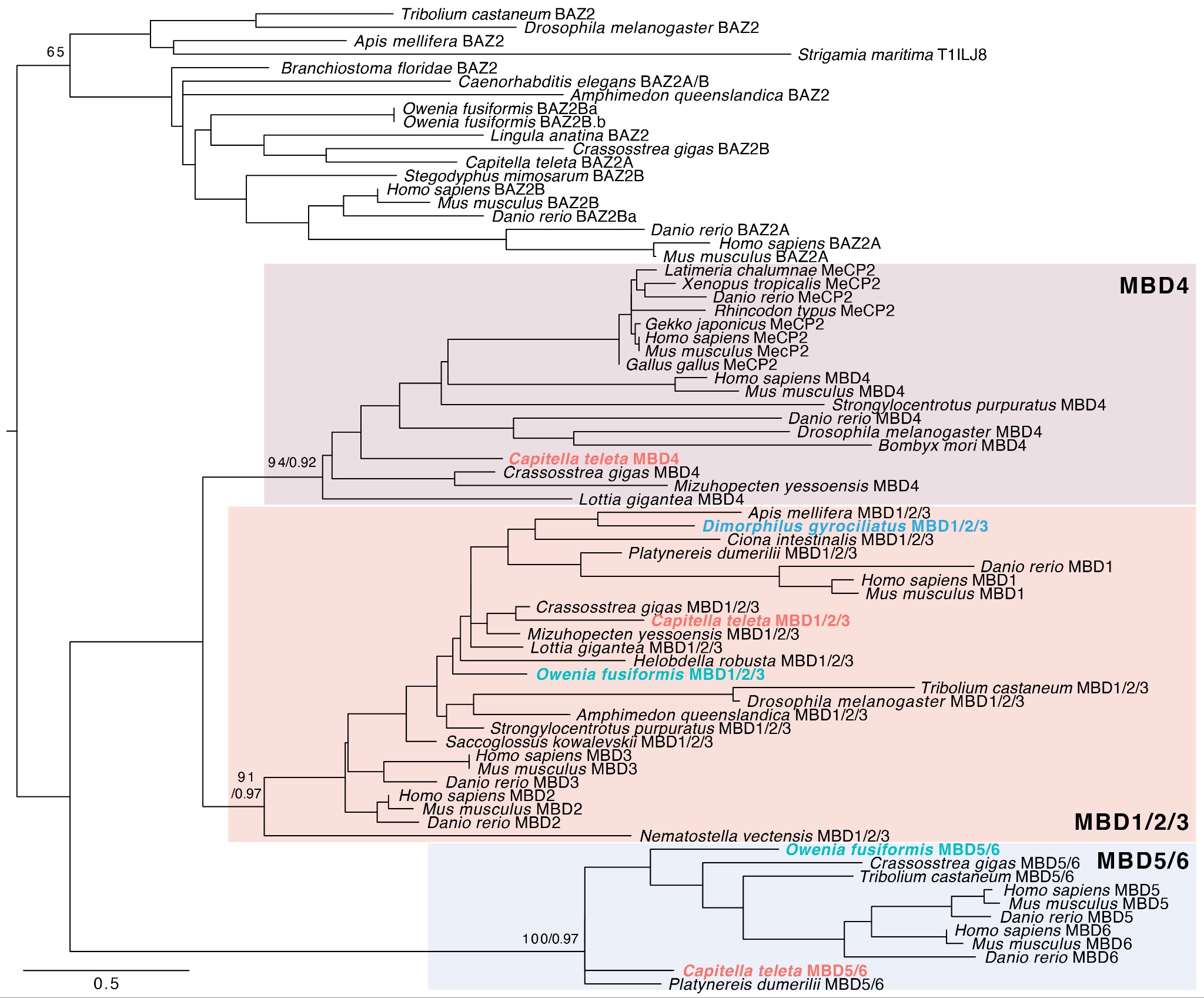

**Figure S2 – Gene orthology assignment of MBD genes.** Orthology assignment of methyl-CpG-binding domain (MBD) proteins using the Bromodomain Adjacent to Zinc Finger Domain 2 (BAZ2) protein as outgroup. The tree topology is based on maximum likelihood reconstruction and node supports indicate both bootstrap values (from 0 to 100) and posterior probabilities (from 0 to 1) at key nodes. Boxes indicate the MBD sub-families and the scale bar represents the number of amino acids substitutions per site alongside the branches.

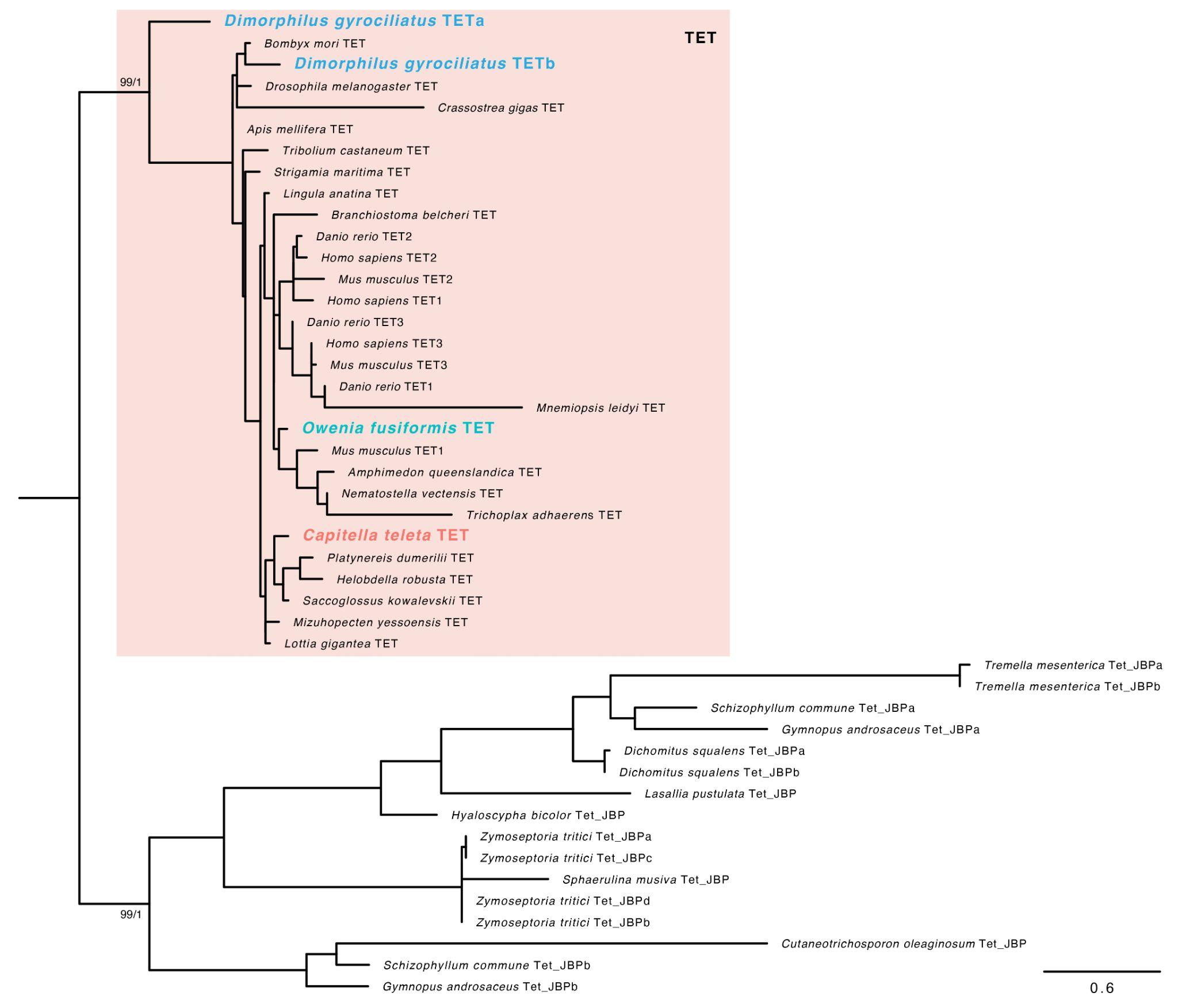

**Figure S3 – Gene orthology assignment of TET genes.** Orthology assignment of ten-eleven translocation (TET) methylcytosine dioxygenases using the TET-domain containing fungal proteins as outgroup. The tree topology is based on maximum likelihood reconstruction and node supports indicate both bootstrap values (from 0 to 100) and posterior probabilities (from 0 to 1) at key nodes. Boxes indicate animal TETs and the scale bar represents the number of amino acids substitutions per site alongside the branches.

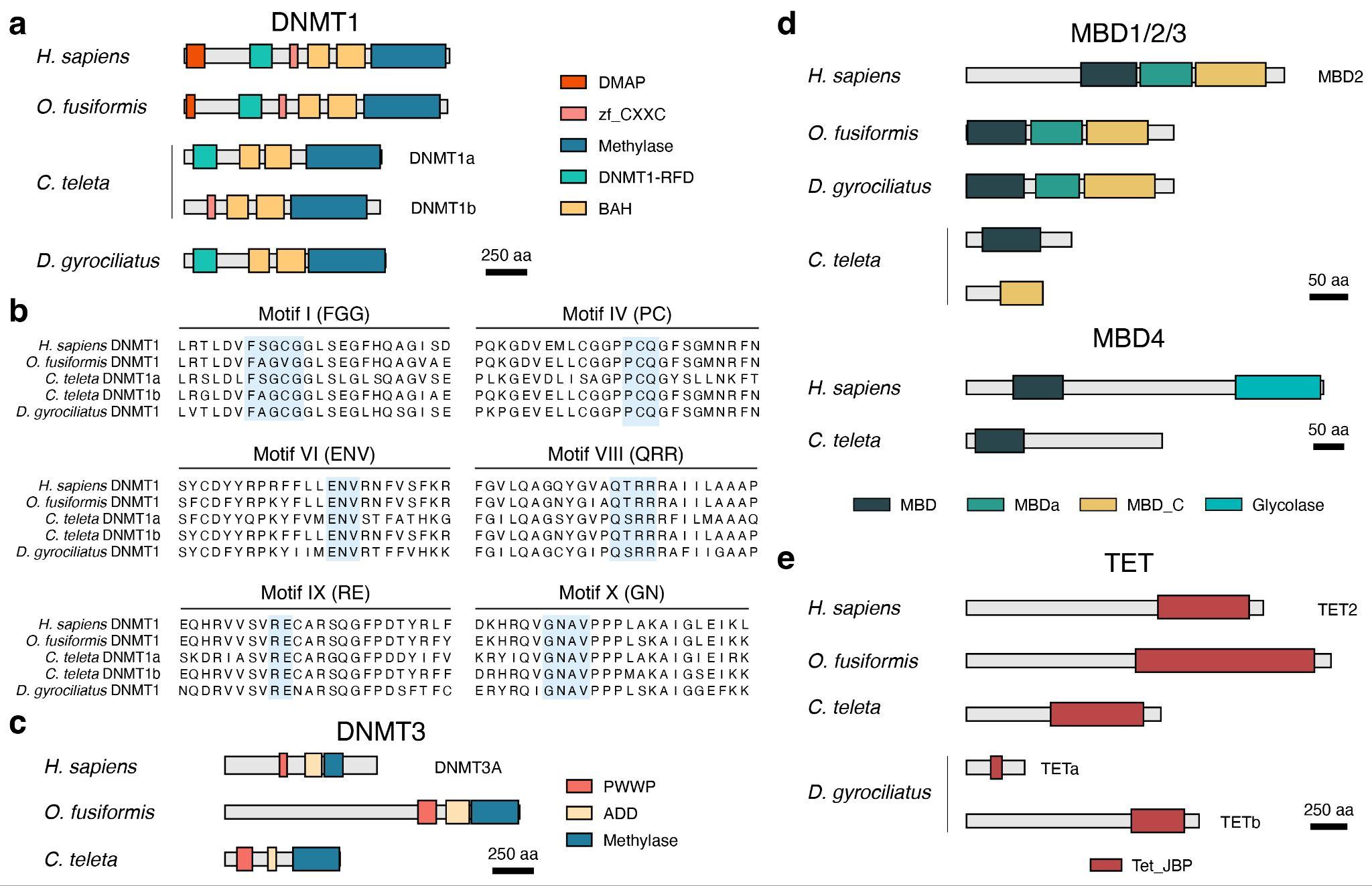

**Figure S4 – The domain architecture of the annelid DNA methylation toolkit.** (**a**) Schematic drawings of the domain composition of DNMT1 genes in the three focal annelid species compared with the human orthologue. (**b**) Multiple protein alignments of the motifs of the DNA methyltransferase domain with catalytic activity comparing annelid (*O. fusiformis*, *C. teleta* and *D. gyrociliatus*) and human sequences. *Dimorphilus gyrociliatus* has conserved residues in these motifs and, thus, a potentially active domain. (**c**–**e**) Schematic drawings of the domain composition of DNMT3 (**c**), MBDs (**d**) and TET (**e**) genes in the three focal annelid species compared with the human orthologue. In (**a**, **c**–**e**), drawings are to scale.

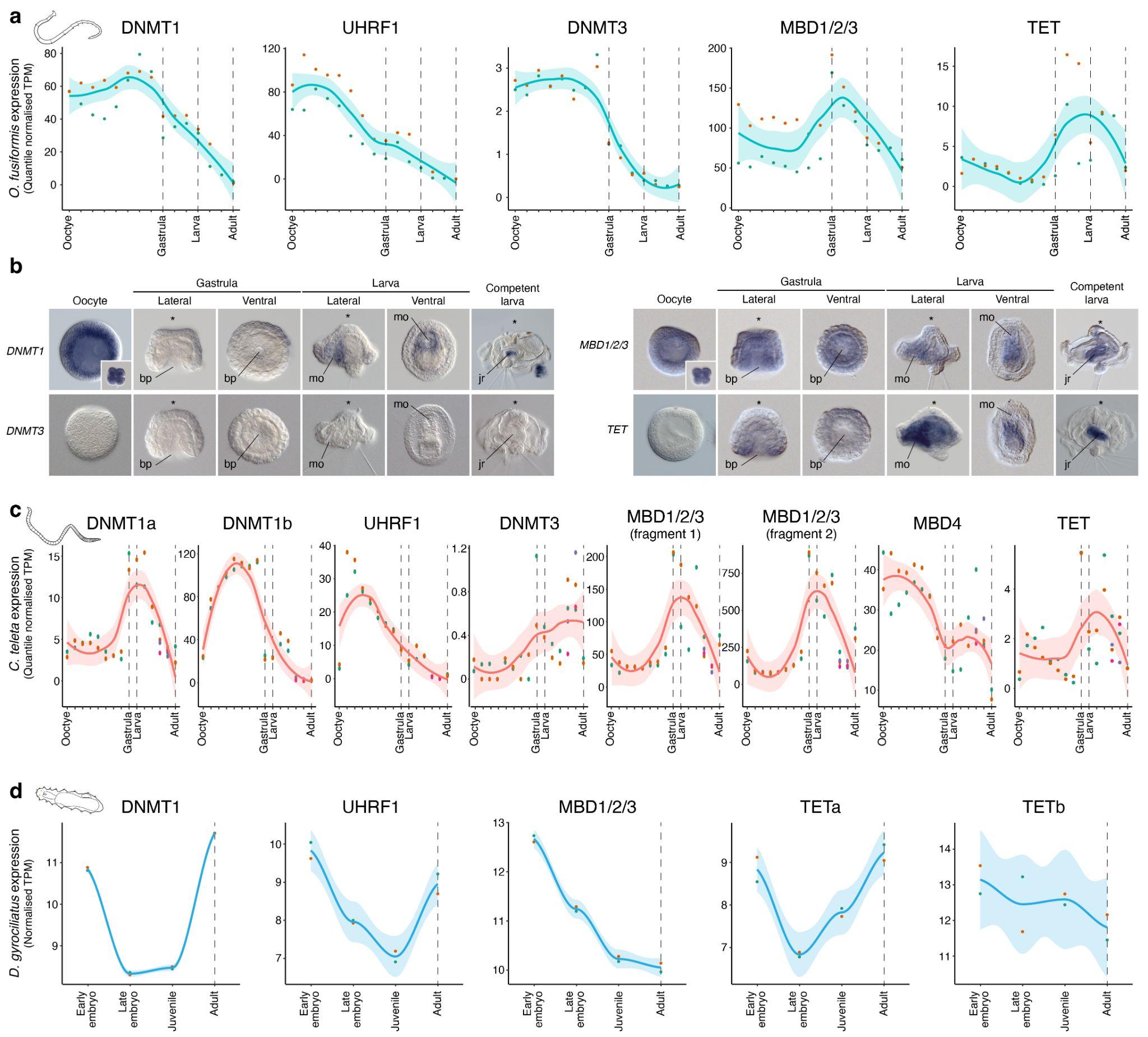

**Figure S5 – The temporal and spatial expression dynamics of the DNA methylation toolkit in Annelida.** (**a**) Line plots depicting the temporal expression dynamics from the active oocyte to the adult stage for each gene of the DNA methylation toolkit in the annelid *O. fusiformis*. (**b**) Photographs of whole-mount in situ hybridisation of *DNMT1*, *DNMT3*, *MBD1/2/3* and *TET* genes during the embryogenesis of *O. fusiformis*. *DNMT1* is strongly expressed in the oocyte and cleavage stages. Its expression decays during mid embryogenesis and it is detected in the ventral side of the early larva and juvenile rudiment of the competent larva. *DNMT3* is expressed at very low levels (**a**) and undetected by in situ hybridisation. *MBD1/2/3* is expressed broadly at all stages of embryogenesis and largely concentrated in the ventral side of the early larva and juvenile rudiment at the competent stage. *TET* is not detected during early embryogenesis and expressed in the gut and ventral side of the early larva and juvenile rudiment of the competent larva. (**c**, **d**) Line plots depicting the temporal expression dynamics from the active oocyte to the adult stage in *C. teleta* (**c**) and from early embryogenesis to the female adult in *D. gyrociliatus* (**d**) for each gene of their respective DNA methylation toolkits. In (**a**, **c**, **d**) the stages sampled for genome-wide methylomes are indicated with dotted vertical lines.

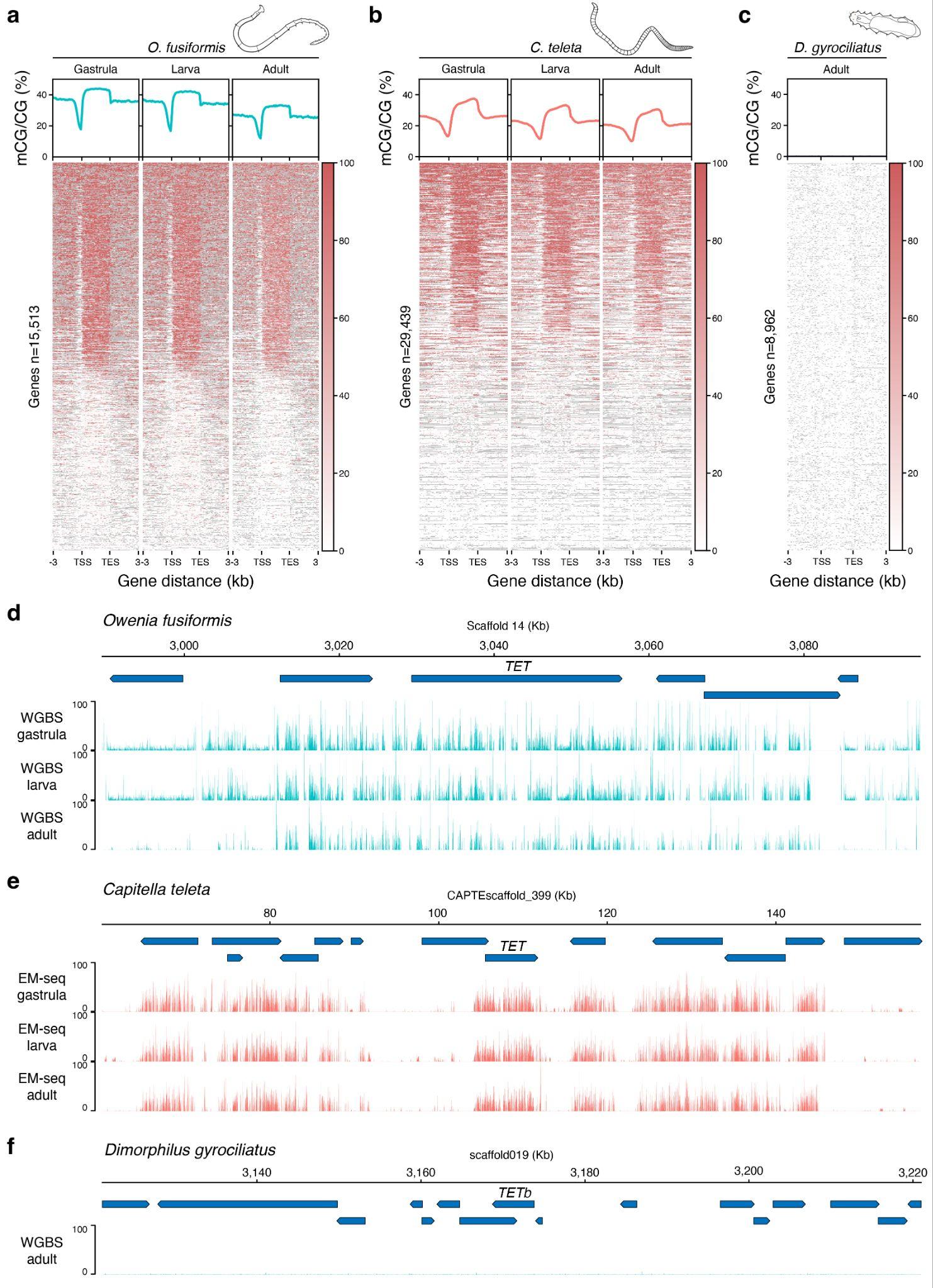

**Figure S6 – The dynamic methylation landscape in Annelida.** (**a**–**c**) Metagene profiles (top) and heatmaps (bottom) of 5mC levels three kilobases upstream and three kilobases downstream of the gene bodies of *O. fusiformis* (**a**), *C. teleta* (**b**) and *D. gyrociliatus* (**c**) in all sampled stages. (**d**–**f**) Genome browser views showing 5mC levels (binning size 75 bases) around the *TET* locus in the three focal annelid taxa in all sampled stages. Transcriptional units are represented by blue boxes ending in an arrowhead that marks the direction of transcription.

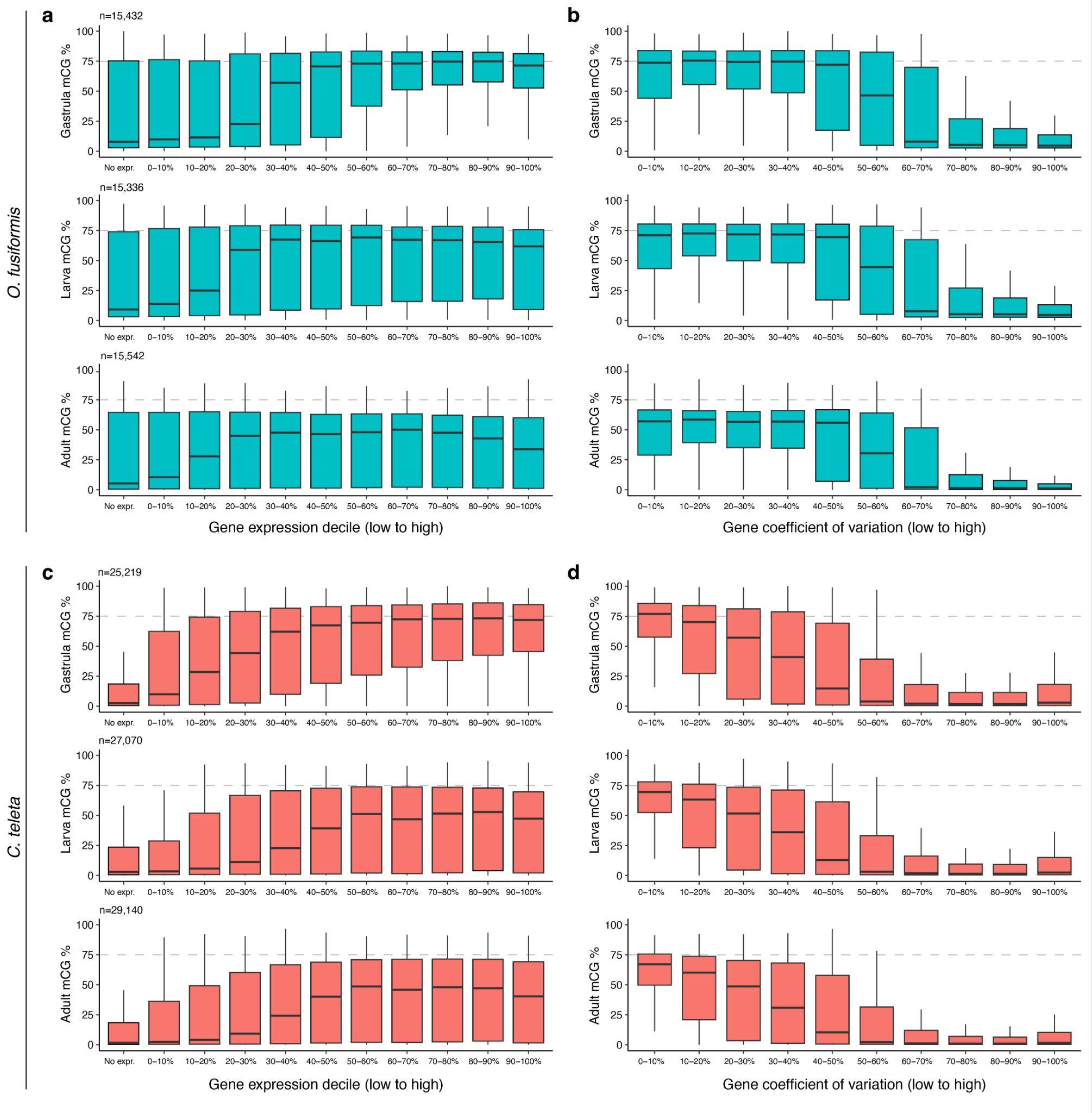

**Figure S7 – DNA methylation and transcriptional dynamics.** (**a**–**d**) Box plots depicting 5mC levels in gene bodies at gastrula, larval and adult stages of *O. fusiformis* and *C. teleta* according to gene expression (**a**, **c**) and gene coefficient of variation (**b**, **d**) deciles. In both species, highly expressed and more stable genes show higher 5mC levels. Notably, gene body methylation levels of the highest express genes decrease as the life cycle progresses in *O. fusiformis* and *C. teleta*.

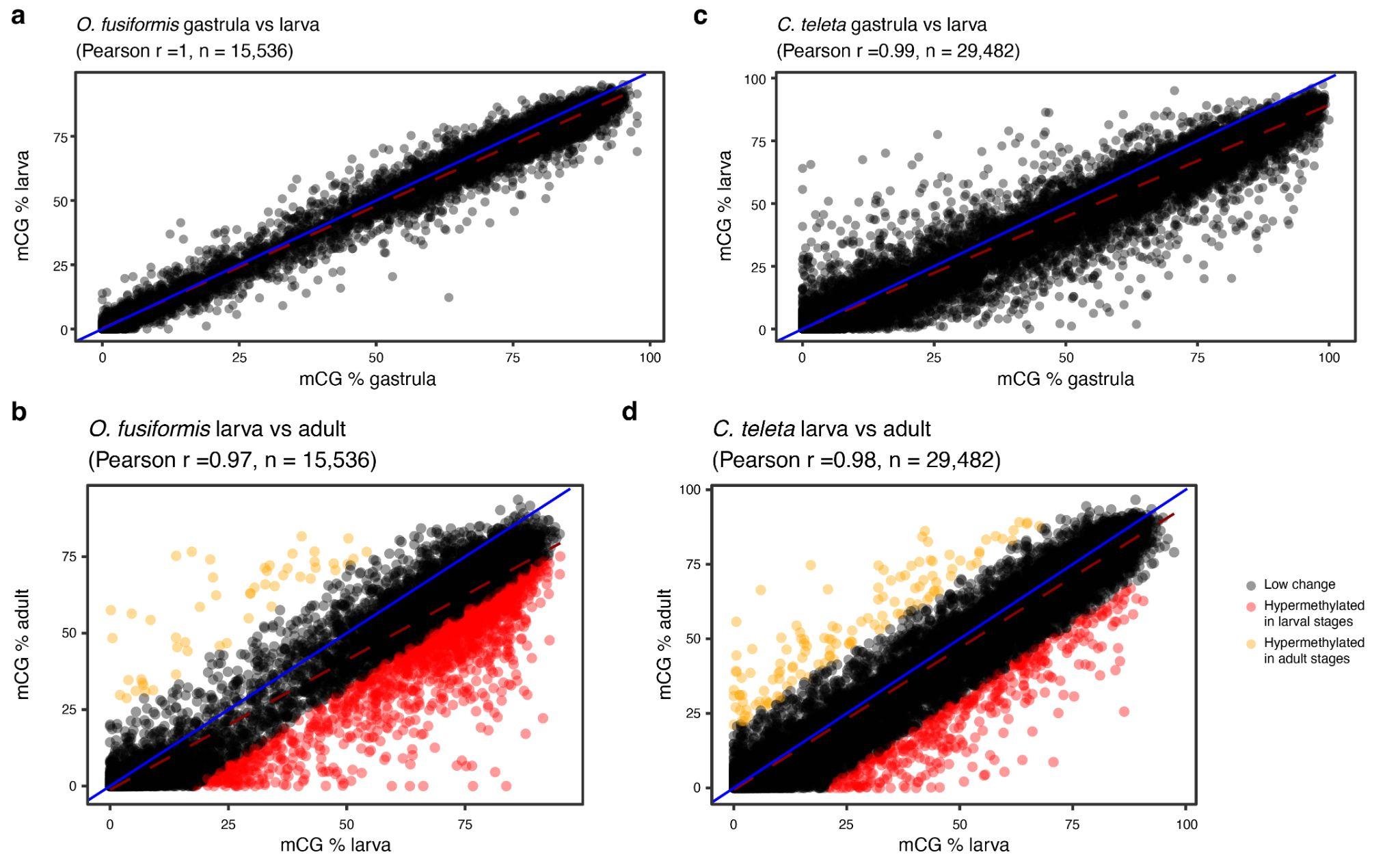

**Figure S8 – Changes in gene body methylation during the annelid life cycle.** (**a**–**d**) Scatter plots of gene body methylation (GbM) levels between consecutive life stages in *O. fusiformis* (**a**, **b**) and *C. teleta* (**c**, **d**). Genes whose GbM changes more than 20% in their methylation status are coloured in red (hypermethylated in the larval stage) and yellow (hypermethylated in the adult stage). A blue line indicates the theoretic diagonal if values were identical across stages, and the dashed red line indicates the correlation slope.

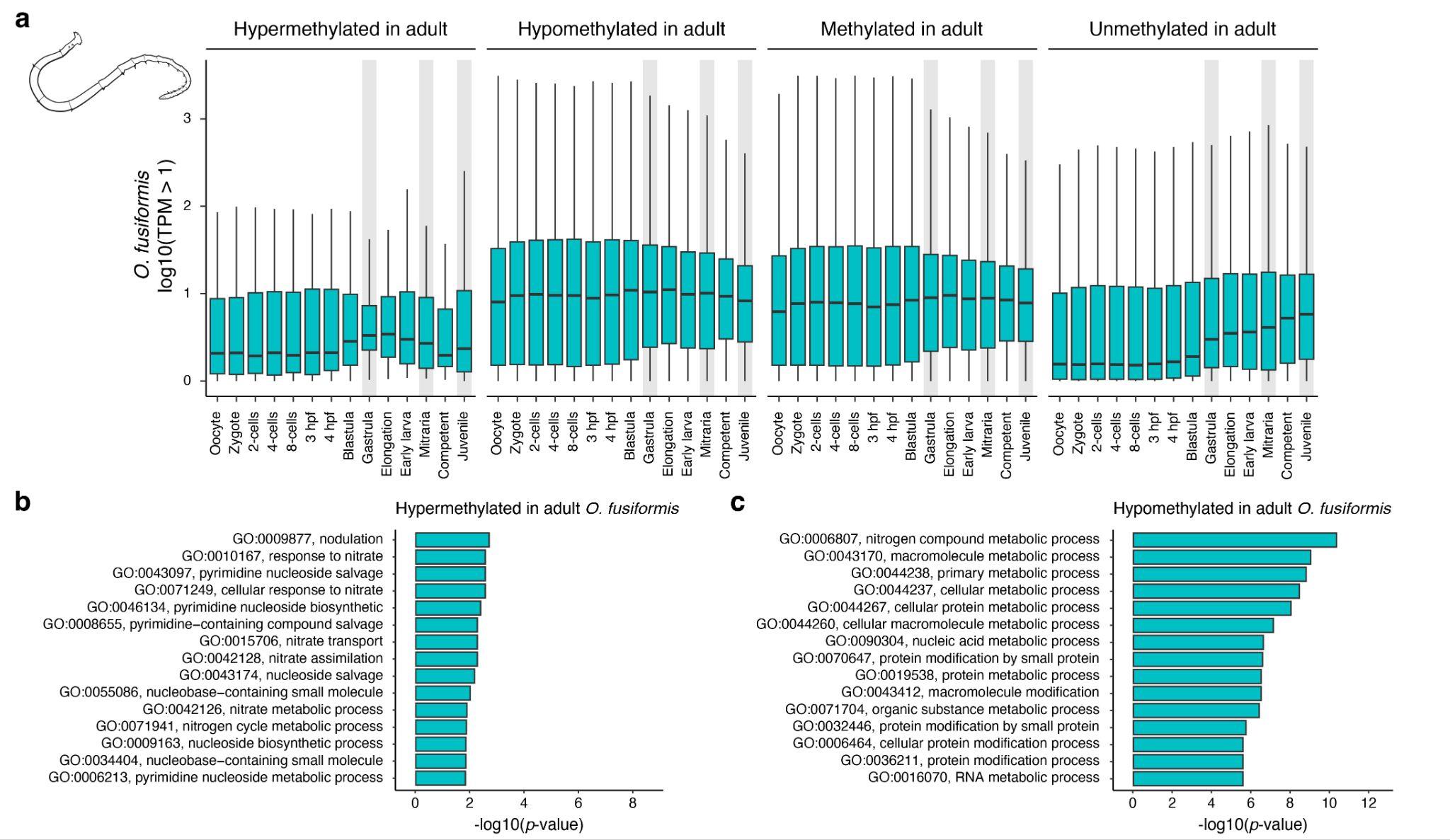

**Figure S9 – The transcriptional dynamics of genes according to their 5mC levels in *O. fusiformis*.** (**a**) Box plots of expression levels for genes that are, from left to right, hypermethylated and hypomethylated in the adult compared to the larval stage, and methylated and unmethylated in the adult of *O. fusiformis*. Grey bars highlight the stages sampled for genome-wide methylomes. (**b**, **c**) Bar plots of Gene Ontology terms enriched in hypermethylated and hypomethylated genes in *O. fusiformis* adults compared to the larval stage.

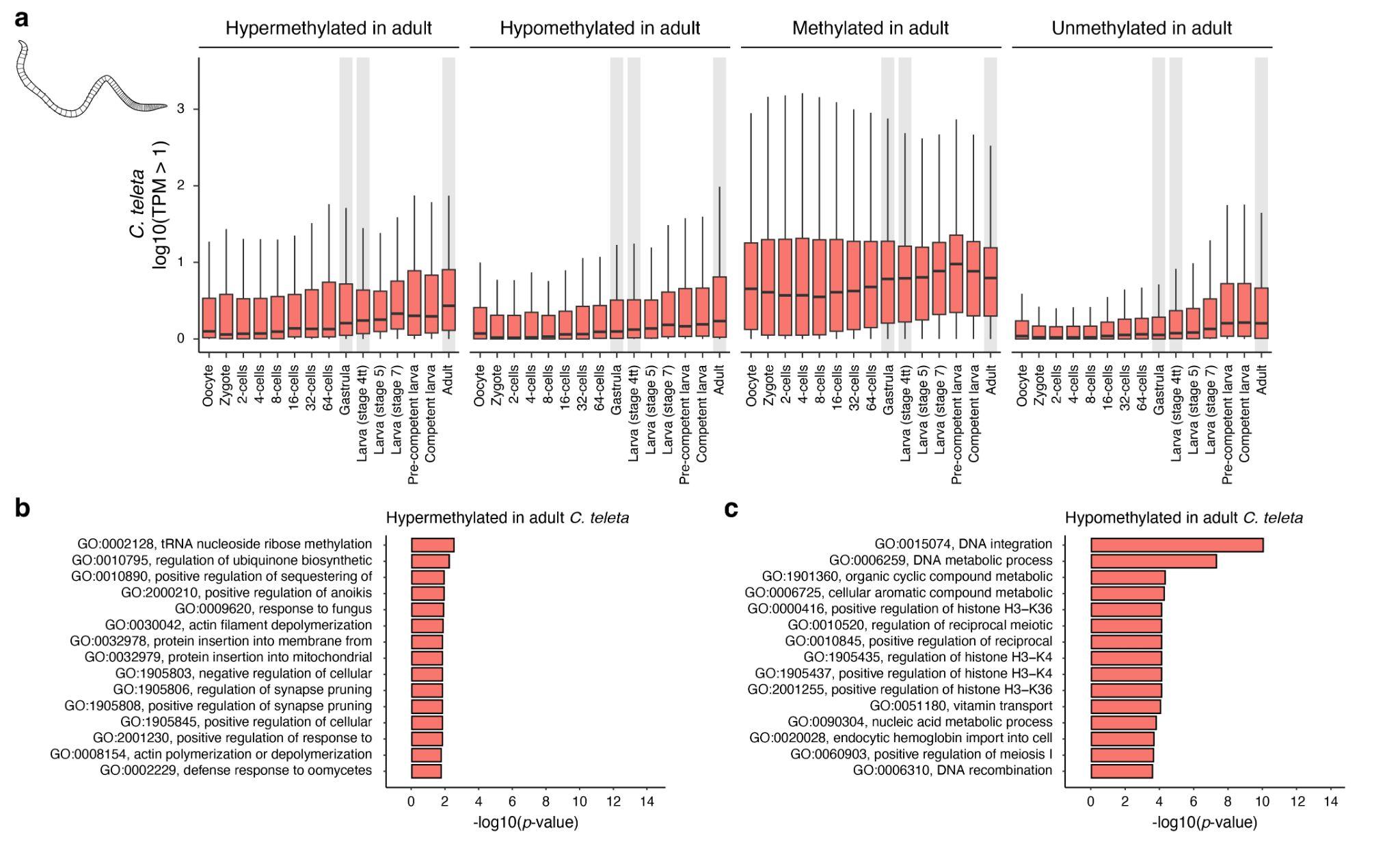

**Figure S10 – The transcriptional dynamics of genes according to their 5mC levels in *C. teleta*.** (**a**) Box plots of expression levels for genes that are, from left to right, hypermethylated and hypomethylated in the adult compared to the larval stage, and methylated and unmethylated in the adult of *C. teleta*. Grey bars highlight the stages sampled for genome-wide methylomes. (**b**, **c**) Bar plots of Gene Ontology terms enriched in hypermethylated and hypomethylated genes in *C. teleta* adults compared to the larval stage.

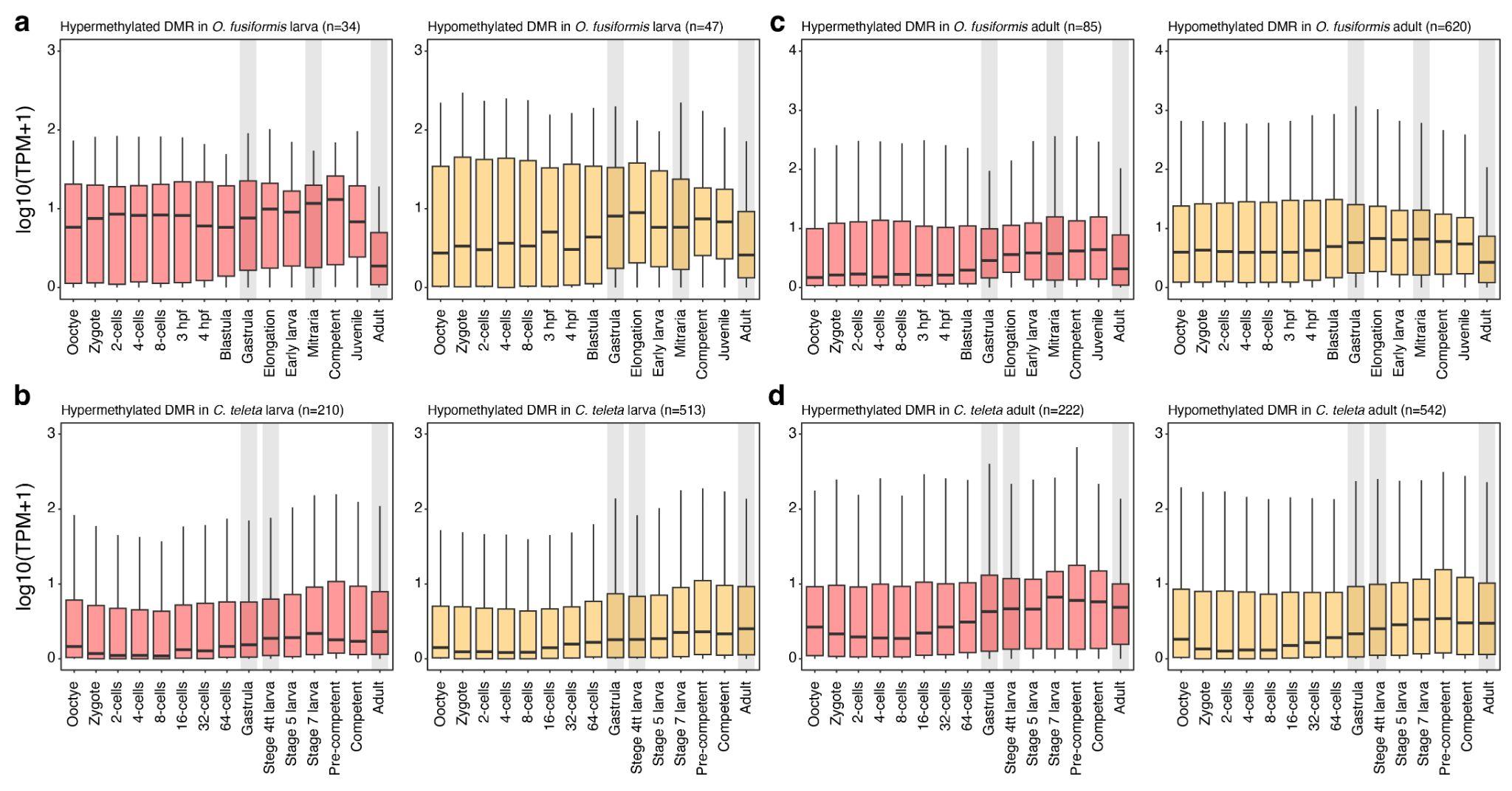

**Figure S11 – The expression dynamics of genes with differentially methylated regions.** (**a**–**d**) Box plots of expression levels from ooctye to adult stages for genes with a hypermethylated (red background) or hypomethylated (yellow background) differentially methylated region (DMR) in its promoter at the larva (**a**) and adult (**b**) stages of *O. fusiformis*, and larva (**c**) and adult (**d**) stages of *C. teleta*. Grey bars highlight the stages sampled for genome-wide methylomes.

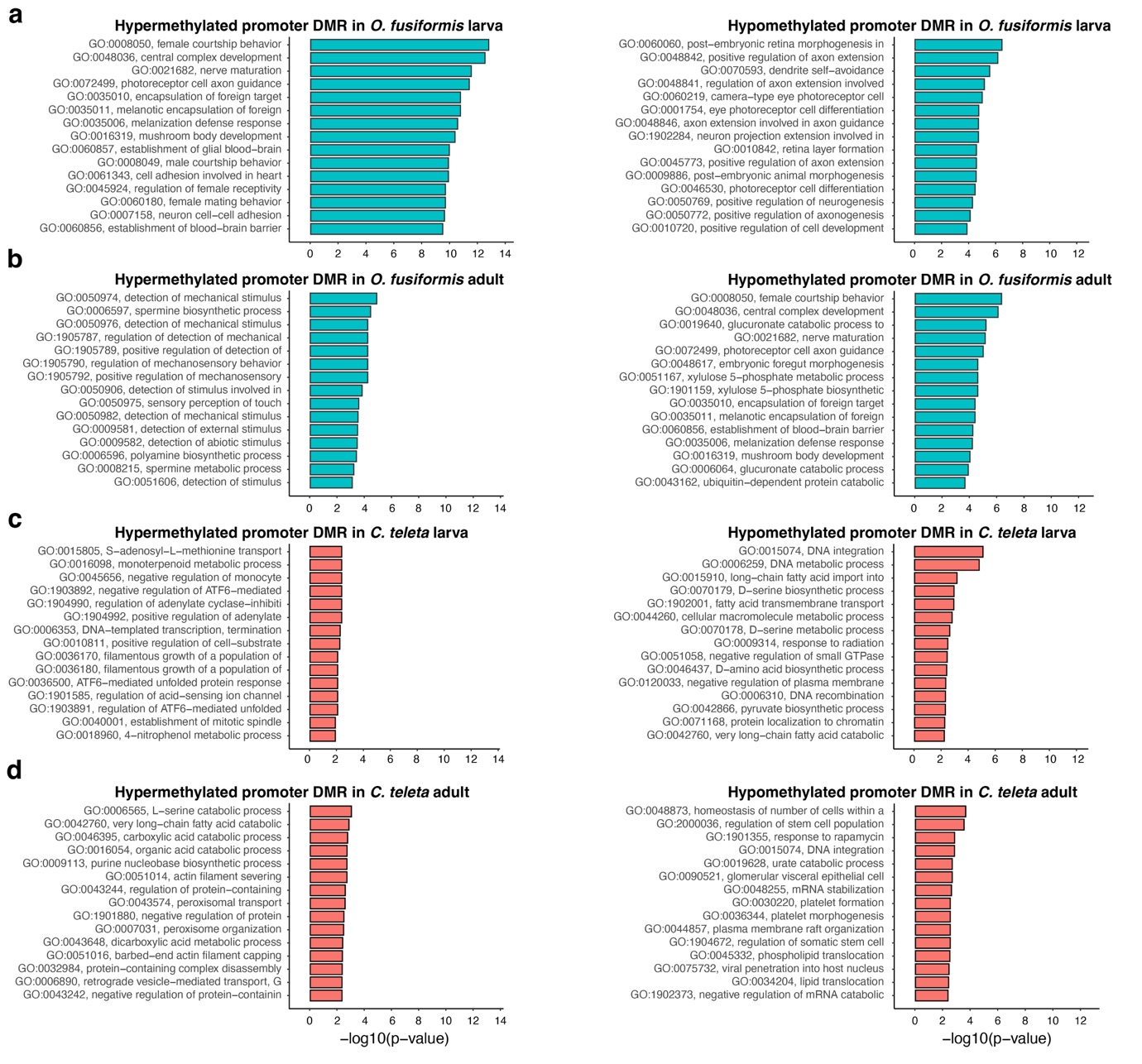

**Figure S12 – The genes experiencing changes in DNA methylation in their promoters during the life cycle in Annelida.** (**a**–**d**) Bar plots of Gene Ontology terms enriched in genes that exhibit a differential methylated region (DMR) in their promoters in the larva (**a**) and adult (**b**) of *O. fusiformis*, and larva (**c**) and adult (**d**) of *C. teleta*.

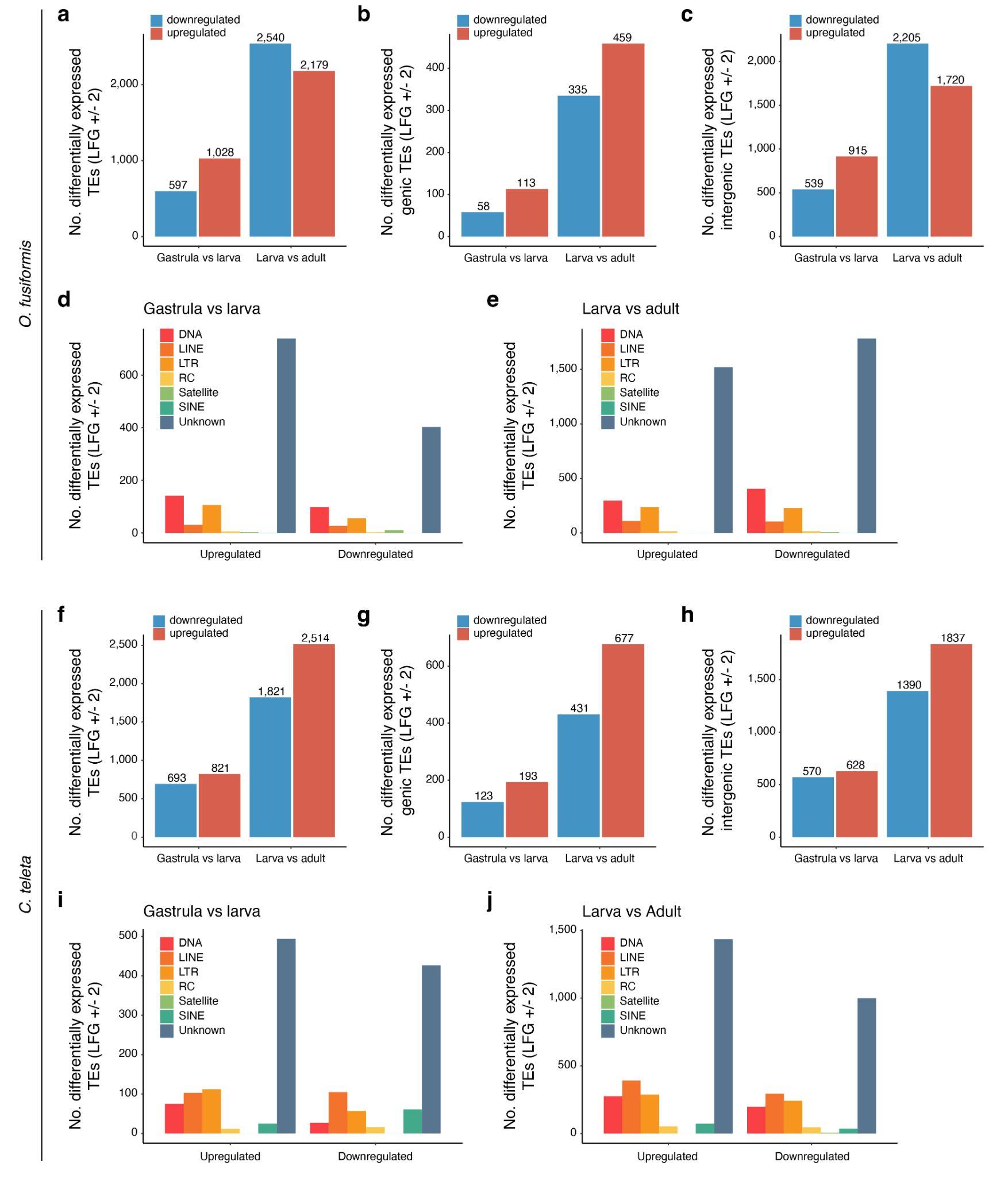

**Figure S13 – The expression dynamics of transposable elements in *O. fusiformis* and *C. teleta*.** (**a**) Bar plots of upregulated and downregulated transposable elements (TE) between the three consecutive life stages sampled for genome-wide methylomes in *O. fusiformis*. (**b**–**e**) Bar plots of upregulated and downregulated TEs between the three consecutive life stages of *O. fusiformis* depending on whether they are within a gene body (genic; **b**) or not (intergenic; **c**) and the TE class (**d**, **e**). (**f**) Bar plots of upregulated and downregulated transposable elements (TE) between the three consecutive life stages sampled for genome-wide methylomes in *C. teleta*. (**g**–**j**) Bar plots of upregulated and downregulated TEs between the three consecutive life stages of *C. teleta* depending on whether they are within a gene body (genic; **g**) or not (intergenic; **h**) and the TE class (**i**, **j**)

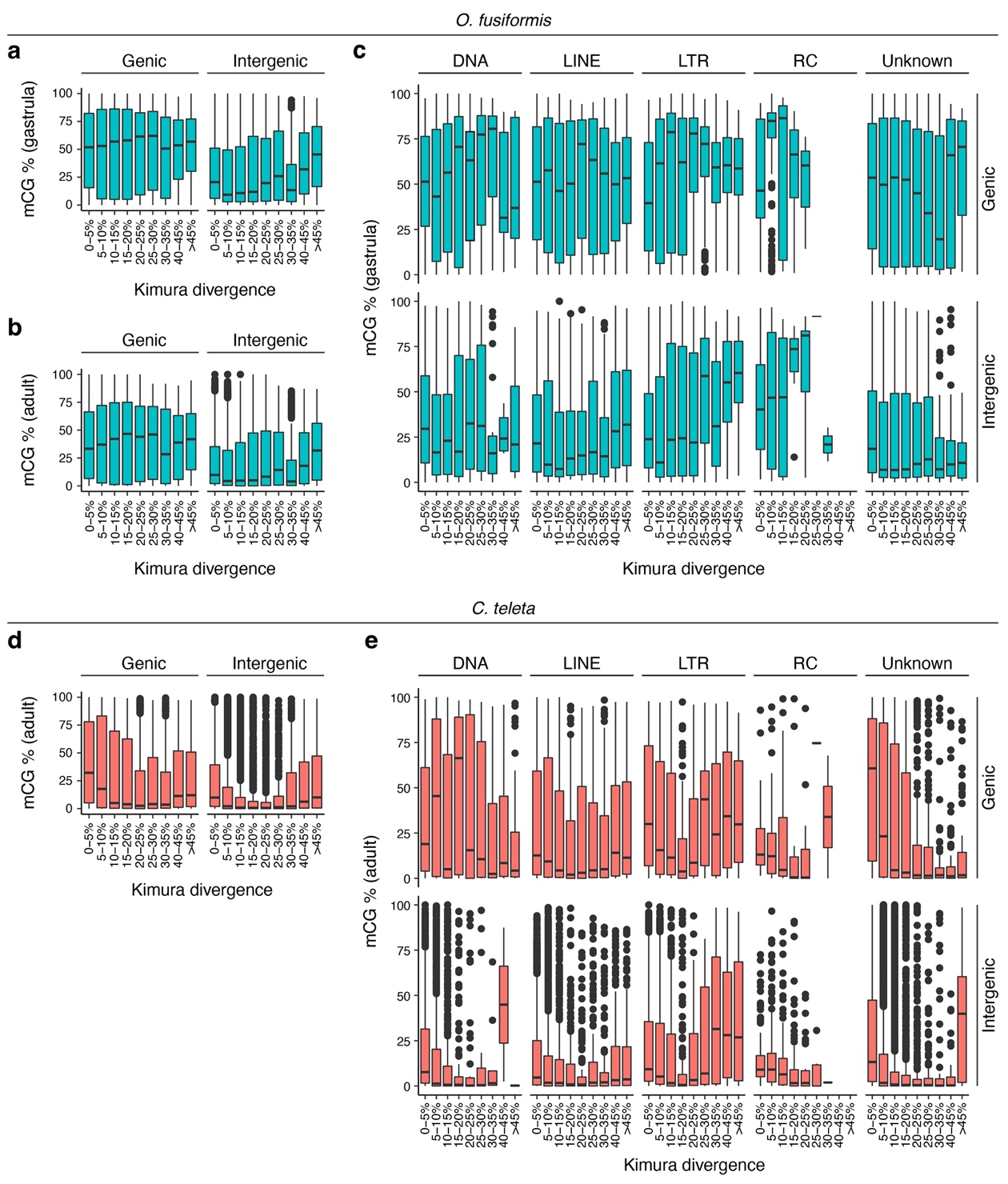

**Figure S14 – 5mC levels and transposable elements in *O. fusiformis* and *C. teleta*.** (**a**–**e**) Box plots of 5mC levels in transposable elements according to their age estimated from the Kimura divergence value in *O. fusiformis* (green background) and *C. teleta* (red background). In (**a**, **b**, **d**), transposable elements are subdivided depending on whether they are within a gene body (genic) or not (intergenic). In (**c**) and (**e**), transposable elements are further subdivided according to their class.

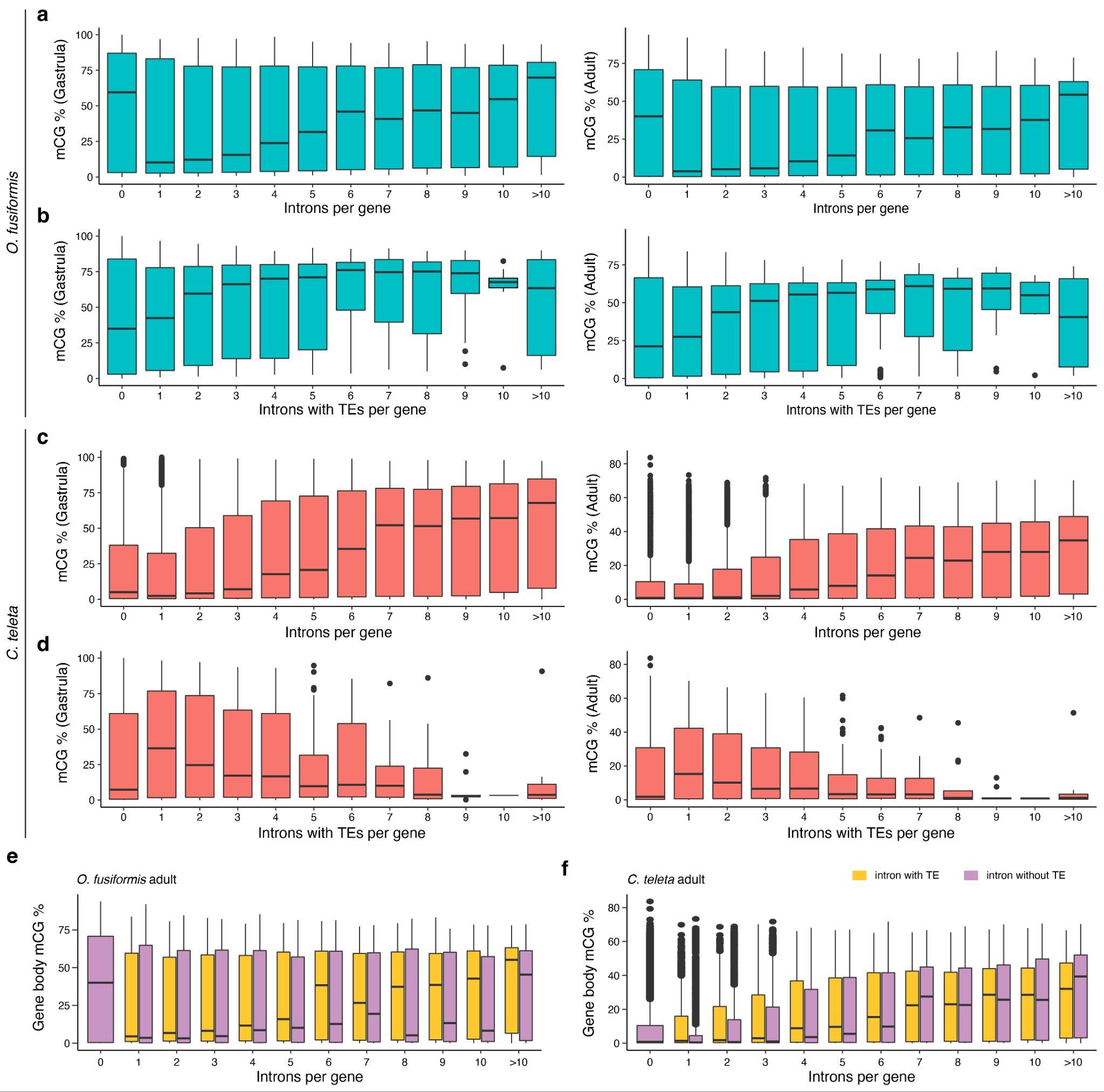

**Figure S15 – The impact of transposable elements in gene body methylation.** (**a**–**d**) Box plots depicting 5mC levels in gene bodies at gastrula (**a**, **c**) and adult (**b**, **d**) stages of *O. fusiformis* (green background) and *C. teleta* (red background) according to the number of introns (**a**, **c**) and number of introns with a transposable element (TE) (**c**, **d**). (**e**, **f**) Gene body methylation levels according to the number of introns and the presence of an intronic TE in *O. fusiformis* (**e**) and *C. teleta* (**f**). Genes with intronic TEs (yellow) have higher gene body methylation levels compared to genes without intronic TEs (violet) in *O. fusiformis* but not *C. teleta*.

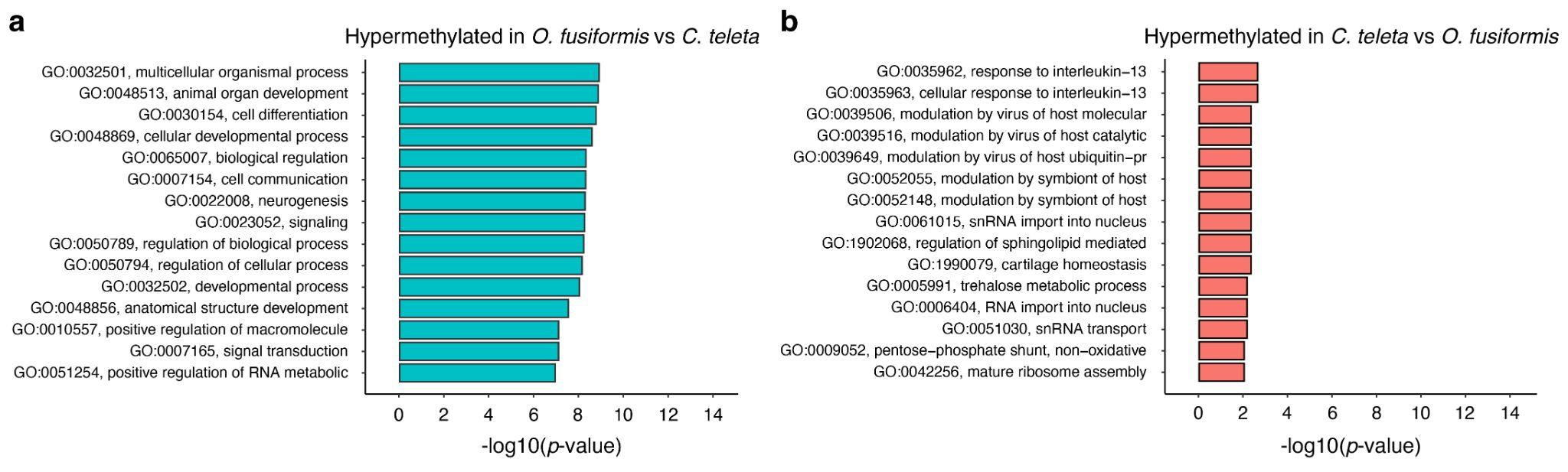

**Figure S16 – The genes with interspecific differences in gene body methylation in Annelida.** (**a**, **b**) Bar plots of Gene Ontology terms enriched in genes that hypermethylated in *O. fusiformis* compared to *C. teleta* (**a**) and hypermethylated in *C. teleta* compared to *O. fusiformis* (**b**).

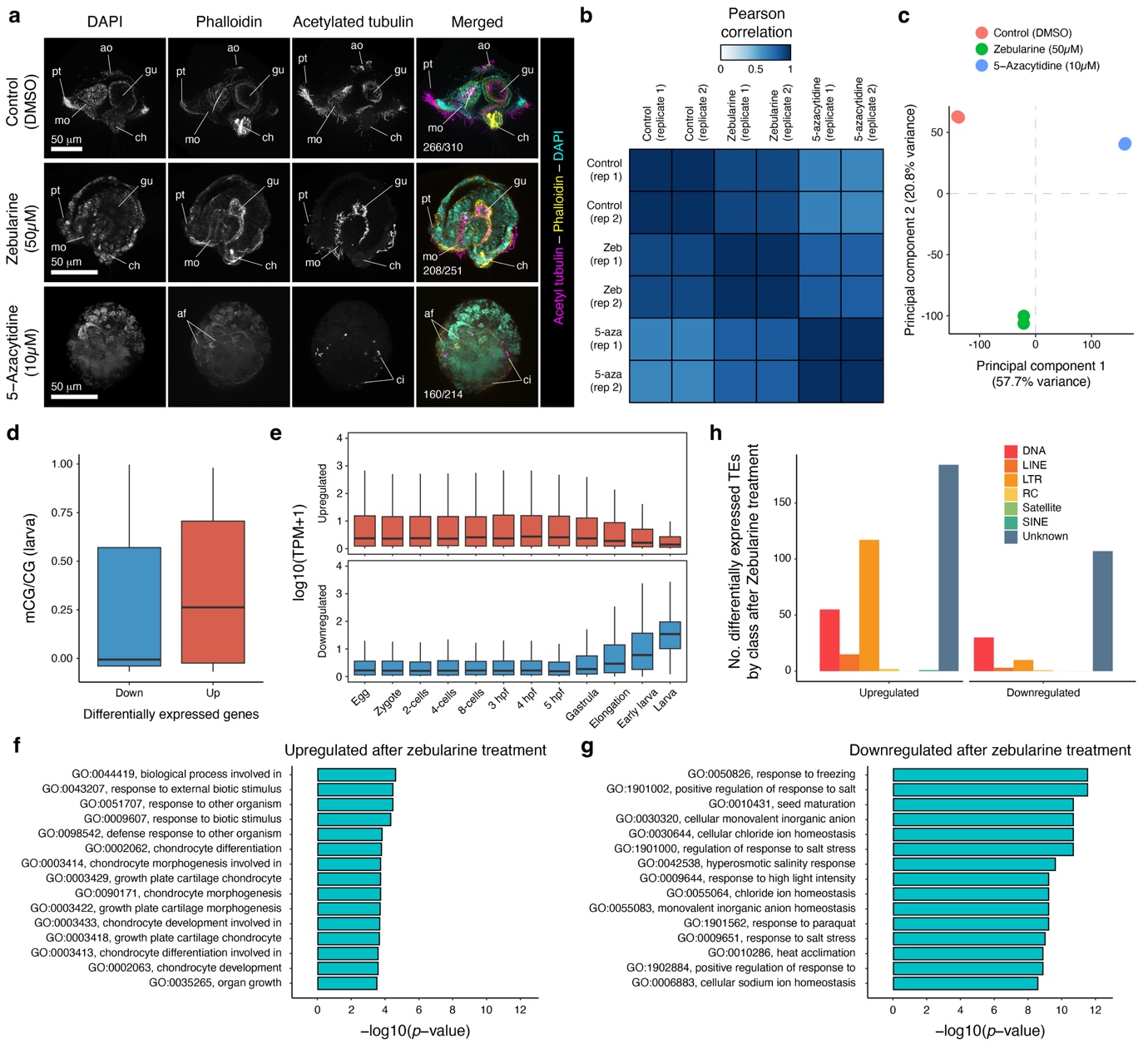

**Figure S17 – The impact of DNA methylation in the early embryogenesis of *O. fusiformis*.** (**a**) Z-projections of confocal stacks of zebularine- and 5-azacytidine-treated and DMSO-control embryos fixed at the early larval stage and stained for acetylated tubulin (magenta), actin (yellow) and nuclei (cyan). Zebularine-treated embryos fail to undergo normal organogenesis in *O. fusiformis*, while 5-azacytidine prevents gastrulation. (**b**) Heatmap of Pearson correlation coefficients and (**c**) principal component analysis of RNA-seq samples of treated and control conditions. (**d**) Box plot of gene body methylation levels at the larval stage of upregulated and downregulated genes after zebularine treatment. (**e**) Bar plots of differentially expressed transposable elements (TEs) by class after zebularine treatment. (**f**, **g**) Bar plots depicting the enrichment of Gene Ontology terms in the gene set that is upregulated (**f**) and downregulated (**g**) after zebularine treatment in *O. fusiformis*.

**Supplementary Table S1. Datasets produced in this study.**

| **Species** | **Stage** | **Sequencing type** | **Depth*** | **Replicates** |
| --- | --- | --- | --- | --- |
| *O. fusiformis* | Gastrula | WGBS | 180.2M | 1 |
|  | Larva | WGBS | 159.4M | 1 |
|  | Adult | WGBS | 255.8M | 1 |
|  | 1% DMSO | EM-seq | 5.6M; 4.2M | 2 |
|  | 50 µM Zebularine | EM-seq | 5.1M; 5.7M | 2 |
|  | 10 µM 5-azacytidine | EM-seq | 6.7M; 4.6M | 2 |
|  | Adult | RNA-seq | 22.7–23.4M | 2 |
|  | 1% DMSO | RNA-seq | 31.8–41.3M | 2 |
|  | 50 µM Zebularine | RNA-seq | 30.8–42.5M | 2 |
|  | 10 µM 5-azacytidine | RNA-seq | 35.6–34.8M | 2 |
| *C. teleta* | Gastrula | EM-seq | 71.3M | 1 |
|  | Larva | EM-seq | 75.1M | 1 |
|  | Adult | EM-seq | 69.3M | 1 |
|  | Competent larva | Nanopore | 446–834 | 2 |
|  | Juvenile | Nanopore | 3,115–4,900 | 2 |
|  | Mature female | Nanopore | 2,535–7,696 | 2 |
|  | Senescent female | Nanopore | 23,959–59,749 | 2 |
|  | Adult | RNA-seq | 27–22.3M | 2 |
| *D. gyrociliatus* | Adult | WGBS | 256.5M | 1 |

*Million paired end reads (RNA-seq, EM-seq and WGBS) and number of reads (Nanopore)

**Supplementary Table S2. Summary of CpG frequency calculations in annelids**

| **Species** | *O. fusiformis* | *C. teleta* | *D. gyrociliatus* |
| --- | --- | --- | --- |
| **CpG dinucleotides** | 8762852 | 9478658 | 1556535 |
| **Cytosine (C)** | 86554427 | 55823136 | 10814336 |
| **Guanine (G)** | 86441988 | 55831156 | 10808663 |
| **Length of genome** | 500139420 | 333283208 | 77897245 |
| **‘N’ nucleotides** | 1020559 | 56624585 | 9614643 |
| **Observed/expected CpGs** | 0.584569 | 0.841396 | 0.909279 |

**Supplementary Table S3. List of drug treatments tested in *O. fusiformis*.** This summarises the concentrations tested across two spawning seasons (2020-2021) of *O. fusiformis*. The asterisk (*) mark indicates the first concentration tested.

| Drug treatments | Concentrations tested (μM) | Results |
| --- | --- | --- |
| Zebularine | 5*, 10, 15, 20, 25, 50 | Dose-dependent effect; morphological defect in larval at 25 μM. Strongest phenotype at 50 µM. |
| 5- Azacytidine | 5*, 10, 15, 20, 25 | Dose-dependent effect; arrested at cell division at 10 μM. |
| Decitabine | 0.1, 1, 2, 5* | Cell death in all concentrations tested. |
